# Microglia contribute to methamphetamine reinforcement and reflect persistent transcriptional and morphological adaptations to the drug

**DOI:** 10.1101/2023.10.19.563168

**Authors:** Samara J. Vilca, Alexander V. Margetts, Isabella Fleites, Claes Wahlestedt, Luis M. Tuesta

**Author notes:** Denotes equal contribution (Samara J. Vilca, Alexander V. Margetts).

## Abstract

Methamphetamine use disorder (MUD) is a chronic, relapsing disease that is characterized by repeated drug use despite negative consequences and for which there are currently no FDA-approved cessation therapeutics. Repeated methamphetamine (METH) use induces long-term gene expression changes in brain regions associated with reward processing and drug-seeking behavior, and recent evidence suggests that methamphetamine-induced neuroinflammation may also shape behavioral and molecular responses to the drug. Microglia, the resident immune cells in the brain, are principal drivers of neuroinflammatory responses and contribute to the pathophysiology of substance use disorders. Here, we investigated transcriptional and morphological changes in dorsal striatal microglia in response to methamphetamine-taking and during methamphetamine abstinence, as well as their functional contribution to drug-taking behavior. We show that methamphetamine self-administration induces transcriptional changes associated with protein folding, mRNA processing, immune signaling, and neurotransmission in dorsal striatal microglia. Importantly, many of these transcriptional changes persist through abstinence, a finding supported by morphological analyses. Functionally, we report that microglial ablation increases methamphetamine-taking, possibly involving neuroimmune and neurotransmitter regulation, and that post-methamphetamine microglial repopulation attenuates drug-seeking following a 21-day period of abstinence. In contrast, microglial depletion during abstinence did not alter methamphetamine-seeking. Taken together, these results suggest that methamphetamine induces both short and long-term changes in dorsal striatal microglia that contribute to altered drug-taking behavior and may provide valuable insights into the pathophysiology of MUD.

**GRAPHICAL ABSTRACT:** 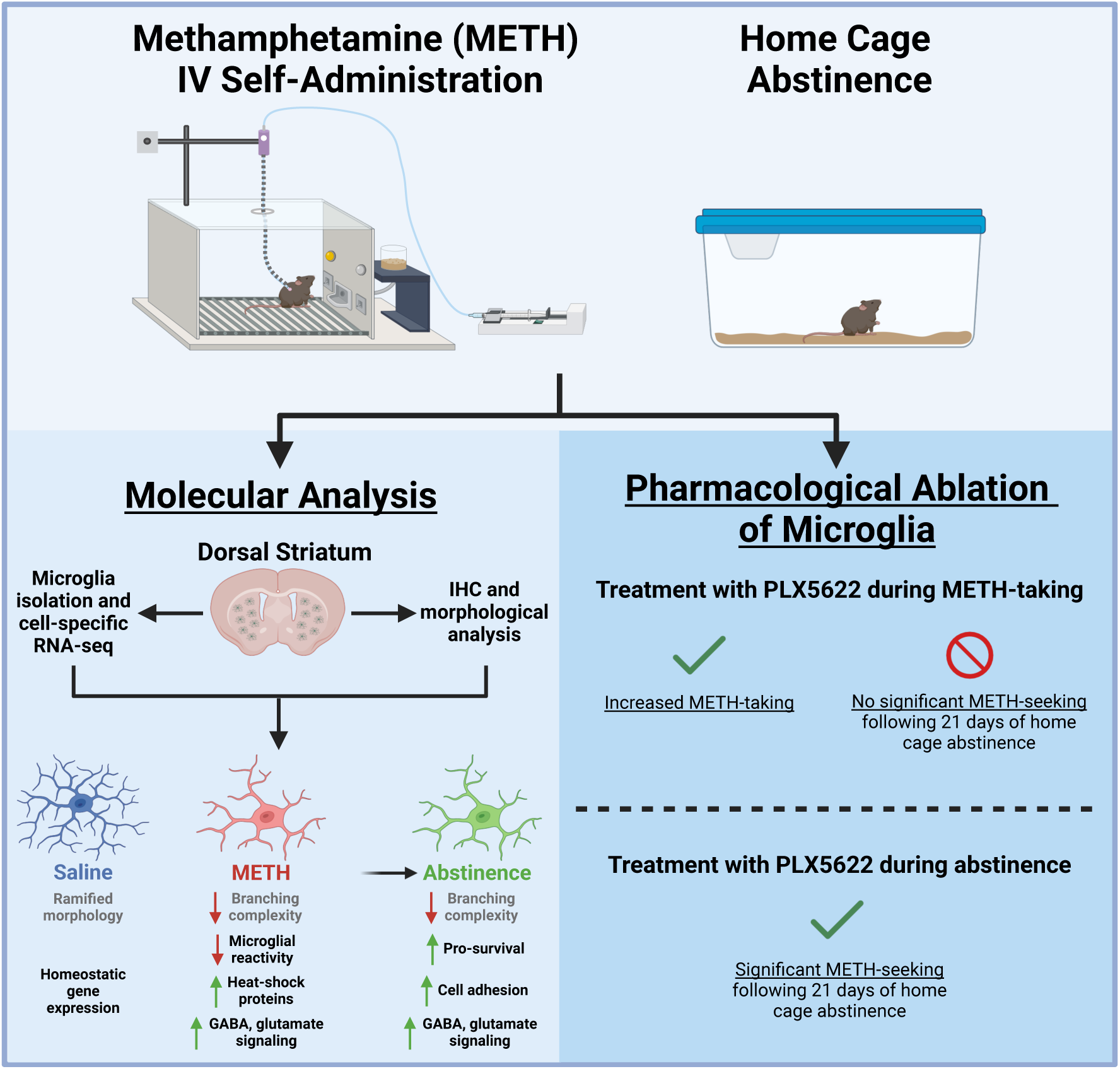

## 1. Introduction

Methamphetamine use disorder (MUD) is a chronic, relapsing disease that is estimated to cost the United States upwards of $24 billion annually (Nicosia, Pacula et al. 2009). Within the past decade, the number of individuals with MUD increased by 37%, while deaths attributed to methamphetamine overdose more than doubled (Centers for Disease Control and Prevention 2021). Methamphetamine has a high potential for addiction due to its potent activation of the brain reward system (Chang, Alicata et al. 2007). Indeed, users can experience intense euphoria, increased energy, and alertness (Cruickshank and Dyer 2009). At higher doses, methamphetamine causes hyperthermia as well as other aversive effects such as arrhythmia, insomnia, paranoia, aggression, and psychosis (Barr, Panenka et al. 2006, Gonzales, Mooney et al. 2010). Methamphetamine reward and reinforcement is attributed to increased dopamine signaling by neurons of the mesocortical, mesolimbic, and nigrostriatal pathways (Everitt and Robbins 2005, Cruickshank and Dyer 2009). Although these neural mechanisms are well characterized, there are currently no FDA approved medications for the treatment of MUD (Karila, Weinstein et al. 2010).

Microglia are the resident immune cell of the central nervous system and have various roles throughout the lifespan including neuronal development and synaptic pruning (Paolicelli, Bolasco et al. 2011), surveillance of the neural environment and circuit formation (Parkhurst, Yang et al. 2013, Wake, Moorhouse et al. 2013), and aging and disease (Keren-Shaul, Spinrad et al. 2017, Salter and Stevens 2017). Accumulating evidence suggests that microglia can respond to methamphetamine exposure (LaVoie, Card et al. 2004, Sekine, Ouchi et al. 2008, Kitamura, Takeichi et al. 2010). For instance, methamphetamine can directly bind several immune receptors expressed in microglia such as TLR4 (Wang, Northcutt et al. 2019) and sigma-1 (Chao, Zhang et al. 2017). Additionally, microglia can respond to methamphetamine-induced neuronal activity and signaling from other glia (LaVoie, Card et al. 2004, Kuhn, Francescutti-Verbeem et al. 2006, Canedo, Portugal et al. 2021). Microglia then release various cytokines which can further amplify the neurotoxic and inflammatory effects of methamphetamine (Krasnova, Justinova et al. 2016). Indeed, long-term activation of microglia may contribute to methamphetamine-related cognitive dysfunction (Sekine, Ouchi et al. 2008, Salamanca, Sorrentino et al. 2014, Liskiewicz, Przybyla et al. 2019). Specifically, striatal microglia have been shown to exhibit a unique transcriptional profiles at baseline (Ayata, Badimon et al. 2018), and after exposure to methamphetamine (Thanos, Kim et al. 2016, Kays and Yamamoto 2019). However, the transcriptional response of microglia to methamphetamine, and particularly prolonged abstinence, have yet to be examined using a clinically relevant animal model of MUD.

Given the evidence suggesting that microglia are actively engaged in the molecular response to methamphetamine, we hypothesized that transcriptional and morphological responses would be more persistent using a clinically relevant model of methamphetamine reinforcement, and that microglia would play a functional role in active methamphetamine-taking. We thus established a model of methamphetamine intravenous self-administration (METH IVSA) in mice as well as a computational pipeline to profile these changes and test the role of microglia in methamphetamine-taking and -seeking following prolonged abstinence.

## 2. Methods

### 2.1. Animals

Male C57BL/6J mice (9 weeks old ∼25-30 g; Jackson Laboratories, Bar Harbor, ME; SN: 000664) were housed in the animal facilities at the University of Miami Miller School of Medicine. Mice were maintained on a 12:12 h light/dark cycle (0600 hours lights on; 1800 hours lights off) and were housed 3 to 5 per cage. Animals were provided with food and water *ad libitum*. Mice representing each experimental group were evenly distributed among testing sessions. All animals were maintained according to the National Institutes of Health guidelines in Association for Assessment and Accreditation of Laboratory Animal Care (AAALAC) -accredited facilities. All experimental protocols were approved by the Institutional Animal Care and Use Committee (IACUC) at the University of Miami Miller School of Medicine. Whenever possible, the experimenter was blind to the experimental and/or treatment group.

### 2.2. Drugs

For self-administration experiments in mice, methamphetamine hydrochloride (NIDA Drug Supply Program, Research Triangle Park, NC, USA) was dissolved in 0.9% sterile saline.

### 2.3. Microglial Depletion

To deplete microglia during METH IVSA, CSF1R inhibitor PLX5622 was formulated in AIN-76A chow (1200 ppm; Research Diets, New Brunswick, NJ, USA). PLX5622 is highly selective to microglia has been shown to ablate nearly all microglia (>97%) when administered for at least 5 days (Spangenberg, Severson et al. 2019) (**Supplementary Fig. 1**). To determine the effect of microglia depletion on methamphetamine-taking, mice were treated with PLX5622 5 days prior to starting METH IVSA, and for the duration of Acquisition and Maintenance. To determine the effect of microglia depletion on methamphetamine-seeking, mice were treated with PLX5622 beginning after the last maintenance session and for the duration of forced home cage abstinence.

### 2.4. Jugular Catheter Surgery

Mice were anesthetized with an isoflurane (1–3%)/oxygen vapor mixture and implanted with an indwelling jugular catheter. Briefly, the catheter consisted of a 6.5-cm length of Silastic tubing fitted to a guide cannula (PlasticsOne, Protech International Inc., Boerne, TX, USA) bent at a curved right angle and encased in dental acrylic resin and silicone. The catheter tubing was passed subcutaneously from the animal’s back toward the right jugular vein, and 1-cm length of the catheter tip was inserted into the vein and anchored with surgical silk sutures. Mice were administered Meloxicam (5 mg/kg) subcutaneously for analgesia prior to start of surgery and 24 hours post-surgery. Catheters were flushed daily with physiological sterile saline solution (0.9% w/v) containing heparin (10–60 USP units/mL) starting 48 hours post-surgery. Animals were allowed 3-5 days to recover from surgery before commencing METH IVSA. Catheter integrity was tested with the ultra-short-acting barbiturate anesthetic Brevital (methohexital sodium, Eli Lilly, Indianapolis, IN, USA).

### 2.5. Operant IV Self-Administration Training

Mice were permitted to self-administer intravenous infusions of either methamphetamine or 0.9% saline during daily 2-hr sessions. Infusions were delivered through Tygon catheter tubing (Braintree Scientific, MA, USA) into the intravenous catheter by a variable speed syringe pump (Med Associates Inc, Fairfax, VT, USA). Self-administration sessions were carried out in operant chambers (Med Associates Inc, Fairfax, VT, USA) containing 2 retractable levers (1 active, 1 inactive) and a yellow cue light located above the active lever which illuminated during the intravenous infusion as well as during the 20-secs post-infusion time out (TO). Completion of the response criteria on the active lever resulted in the delivery of an intravenous infusion (14 µL over 2 sec) of methamphetamine (0.05 mg/kg/infusion) or 0.9% saline. Responses on the inactive lever were recorded but had no scheduled consequences. During Acquisition, mice were trained daily using the following fixed ratio schedule of reinforcement: FR1 for infusions #1-5, FR2 for infusions #6-10, and FR3 for the remainder of the session. Following 5 consecutive days of Acquisition (training), mice were allowed to self-administer methamphetamine or saline at FR3TO20 during 10 consecutive daily 2-hr Maintenance sessions. Animals that did not achieve stable responding (fewer than 7 infusions per 2-hr session) or demonstrated signs of compromised catheter patency, were excluded from analysis.

### 2.6. Forced Home Cage Abstinence and Context-Induced Seeking

Following 15 days of methamphetamine self-administration (Acquisition and Maintenance), mice underwent 21 days of forced home cage abstinence. A context-induced drug-seeking session was then conducted on Day 21, where completion of response criteria resulted in the presentation of the light stimulus previously paired with methamphetamine or saline infusion delivery; however, no reward was delivered. Active lever presses were recorded and interpreted as a measure of relapse.

### 2.7. Microglial Isolation

Immediately after the final self-administration maintenance session (Saline and Maintenance) and following 21 days of forced home cage abstinence (Abstinence), mice were anesthetized with isoflurane and perfused through the ascending aorta with 0.1 M phosphate buffer saline (PBS pH 7.4, Gibco, Waltham, MA, USA) plus heparin (7,500 USP units). Tissues were then immediately dissected and transported in Hibernate A Medium (Gibco) before dissociation. Briefly, tissue was enzymatically and mechanically dissociated, and debris removed using the Adult Brain Dissociation Kit (Miltenyi Biotec, Bergisch Gladbach, Germany). The resulting single cell suspension was incubated with anti-mouse CD11b (i.e., microglia-specific) magnetic MicroBeads (Miltenyi Biotec, #130–093–634) and microglia were positively selected for via column purification (Miltentyi Biotec, #130-042-201). Purity of resulting microglial samples were confirmed by enrichment of microglia-specific genes and depletion of genes associated with macrophages, neurons, astrocytes, and oligodendrocytes (**Supplementary Fig. 2**).

### 2.8. Brain Perfusion and Fixation

Mice were anesthetized with isoflurane and perfused through the ascending aorta with PBS pH 7.4 (Gibco) plus heparin (7,500 USP units), followed by fixation with 4% paraformaldehyde in PBS. Brains were collected and postfixed overnight in 4% paraformaldehyde, then transferred to 30% sucrose with 0.05% sodium azide (S2002, Sigma-Aldrich, St. Louis, MO, USA) in PBS for 72 hrs. All brains were cut into 35 µm coronal sections on a Leica CM1900 cryostat and placed into 12-well plates containing PBS with 0.05% sodium azide at 4°C until processing for immunohistochemistry.

### 2.9. Fluorescence Immunolabeling

Free-floating brain sections were processed for fluorescence immunostaining of dorsal striatal microglia. Sections were rinsed in PBS and then blocked for 1 hour in Blocking Buffer consisting of 10% normal donkey serum (017-000-121, Jackson ImmunoResearch), 0.5% Triton X-100 (T8787, Sigma), and PBS. Thereafter, sections were incubated in primary antibody diluted in Blocking Buffer overnight at 4°C. The following primary antibodies were used: Rabbit Anti-Iba1 (1:1000, Wako Fujifilm, 019-19741), Mouse Anti-NeuN (1:1000, Millipore, MAB377), Mouse Anti-GFAP (GA5) (1:500, Cell Signaling Technology, 3670), Mouse Anti-APC (1:100, Millipore, OP80). On day 2, sections were washed in PBS three times for 5 min each, then incubated with the following secondary antibody: Alexa Fluor 488 Donkey Anti-Rabbit (1:500, A-21206, Invitrogen), Alexa Fluor 568 Donkey Anti-Mouse (1:500, A-10037, Invitrogen). Sections were incubated with secondary antibodies in PBS with 2% normal donkey serum for 2 hours at room temperature in the dark. Next, sections were rinsed in PBS three times for 5 min each and mounted on slides with VECTASHIELD Antifade Mounting Medium with DAPI (Vector Laboratories, H-1200-10) and coverslipped. Fluorescent images were acquired on an ECHO Revolve microscope using 20x and 60x objectives. All antibodies used have been previously validated for the intended applications, as per manufacturer. For all representative images of qualitative data, the immunolabeling experiment was successfully repeated in 4 animals.

### 2.10. Sholl Analysis

Acquired images were converted to 8-bit grayscale and analyzed using FIJI (Schindelin, Arganda-Carreras et al. 2012). Iba1-positive channel was enhanced across the entire image, followed by noise de-speckling. The image was then converted to binary and skeletonized. Microglia morphology was analyzed using FIJI’s Sholl Analysis plugin (Ferreira, Blackman et al. 2014). Briefly, the upper limit for concentric circle placement was set by drawing a radius from the center of the cell soma to the end of the longest branch. Then, the starting radius was set at 5 µm with a step size of 2 µm. Finally, the number of branch interceptions at each of the concentric circles was calculated. Each condition consisted of 40-43 counted microglia (10-12 microglia per animal) from a total of 4 animals. Only microglia located in the dorsal striatum whose soma and processes were completely within the field-of-view and in focus were considered for analysis.

### 2.11. RNA-Sequencing

Isolated dorsal striatal microglia (n = 5-7 per condition) were centrifuged for 5 min at 600xg and resuspended in RLT plus buffer (Qiagen) for extraction and purification of total RNA. RNA input was normalized and NGS libraries were prepared using NEBNext Single Cell/Low Input RNA Library Prep Kit for Illumina (New England BioLabs) according to the manufacturer’s instructions. Paired-end 100 bp sequencing was performed on a NovaSeq6000 sequencer (Illumina). All RNA-seq data used in this study were mapped to the mm10 genome. Prior to mapping, raw RNA-seq datasets were trimmed using Trimgalore (v.0.6.7) and cutadapt (v.1.18). Illumina sequence adaptors were removed and the leading and tailing low-quality base-pairs were trimmed following default parameters. Next, pair-end reads were mapped to the genome using STAR (v.2.7.10a) with the following parameters: –outSAMtype BAM SortedByCoordinate –outSAMunmapped Within – outFilterType BySJout –outSAMattributes NH HI AS NM MD XS –outFilterMultimapNmax 20 – outFilterMismatchNoverLmax 0.3 --quantMode TranscriptomeSAM GeneCounts. The resulting bam files were then passed to StringTie (v.2.1.5) to assemble sequenced alignments into estimated transcript and gene count abundance given the NCBI RefSeq GRCm38 (mm10) transcriptome assembly.

### 2.12. Differential Gene Expression Analysis

The R/Bioconductor DESeq2 package (v.1.38.3) was used to detect the differentially expressed genes in microglia throughout different phases. Following filtering for low count genes and outliers, as determined by DESeq2 and Cook’s distance, and using a false discovery ratio (FDR) correction, only genes with an adjusted *p-value* < .05 were considered as significantly differentially expressed. In the case where biological replicates showed high variability, indicating outliers, a supervised removal of such replicates was conducted, leaving an n = 5-7 per condition for downstream analysis (**Supplementary Fig. 3**).

### 2.13. Functional Enrichment Analysis

The enrichGO function from the R/Bioconductor clusterProfiler package (v.4.6.2) was used to perform gene ontology (GO) enrichment analysis. Only significantly differentially expressed genes with an adjusted *p-value* < .05 and lfcSE ≤ 1.5 were included. Resulting GO terms and pathways with an FDR < .05 were considered after using a custom background from all genes that were expressed after DESeq2 adjustment. The associated GO and pathway enrichment plots were generated using the ggplot2 package. Heatmaps were generated using the R/Bioconductor package pheatmap (v.1.0.12) of regularized log (rlog) transformed normalized counts. All the other plots were generated using the ggplot2 package (v.3.4.2) with labels added using Adobe Illustrator for clarity.

### 2.14. Statistical analyses

Animal sample size was justified by previously published data or preliminary experiments. Data distribution was assumed to be normal. All animals were randomly assigned to treatment groups. For self-administration experiments, animals that did not achieve stable levels of intake (<20% variation in intake across three consecutive days) or that took fewer than 7 methamphetamine infusions on average across sessions were excluded from data analysis. All behavioral and immunohistochemical data were analyzed by Two-way RM ANOVA, One-way ANOVA, or t-tests using GraphPad Prism software (La Jolla, CA). Significant main or interaction effects were followed by appropriate multiple comparisons tests. The criterion for significance was set at < .05. Data are shown as the mean ± SEM.

### 2.15. Code and Data availability

All next generation sequencing files associated with this study as well as the code that was used to pre-process and run differential expression are available online https://avm27.github.io/Methamphetamine_MicroglialRNASequencing_Analysis/.

## 3. Results

### 3.1. Mice acquire and maintain stable methamphetamine-taking, and demonstrate methamphetamine-seeking following forced home cage abstinence

To profile microglial gene expression changes during methamphetamine-taking and -seeking, we first established a model of METH IVSA. Mice underwent METH IVSA and forced home cage abstinence according to the timeline (**Fig. 1A**). Mice self-administered significantly more methamphetamine than saline during Maintenance (**Fig. 1B**) (Two-way RM ANOVA; METH vs Saline, F (1, 19). = 25.75, *p* < .0001). Additionally, mice self-administering methamphetamine displayed robust lever discrimination, while saline-taking mice did not (**Fig. 1C**) (Two-way RM ANOVA; Active vs Inactive Lever, F (3, 38) = 29.49, *p* < .0001; METH vs Saline, F (1, 19) = 17.46, *p* = .0005). Following Abstinence, mice underwent a 2-hr context-induced drug-seeking session (**Fig 1D**). Mice that previously self-administered methamphetamine showed increased active-lever responding (**Fig. 1E**) (Two-way RM ANOVA; METH Maint vs Seek, F (1, 19) = 6.93, *p* = .016; METH vs Saline Seek, F (1, 19) = 11.73, *p* = .003), as well as significant lever discrimination compared to control mice that had self-administered saline (**Fig. 1F**) (Two-way ANOVA; METH Active vs Inactive Lever, F (1, 38) = 13.82, *p* = .0006; METH vs Saline, F (1, 38) = 6.29, *p* = .016). Indeed, in our model of METH IVSA, operant responding was higher for mice infusing methamphetamine than mice infusing saline, supporting the reinforcing properties of methamphetamine. Further, mice exhibited a high degree of lever discrimination when self-administering methamphetamine and showed higher rate of active lever pressing during the drug-seeking session, demonstrating the ability of this model to recapitulate methamphetamine-taking and - seeking behavior.

**Figure 1.**
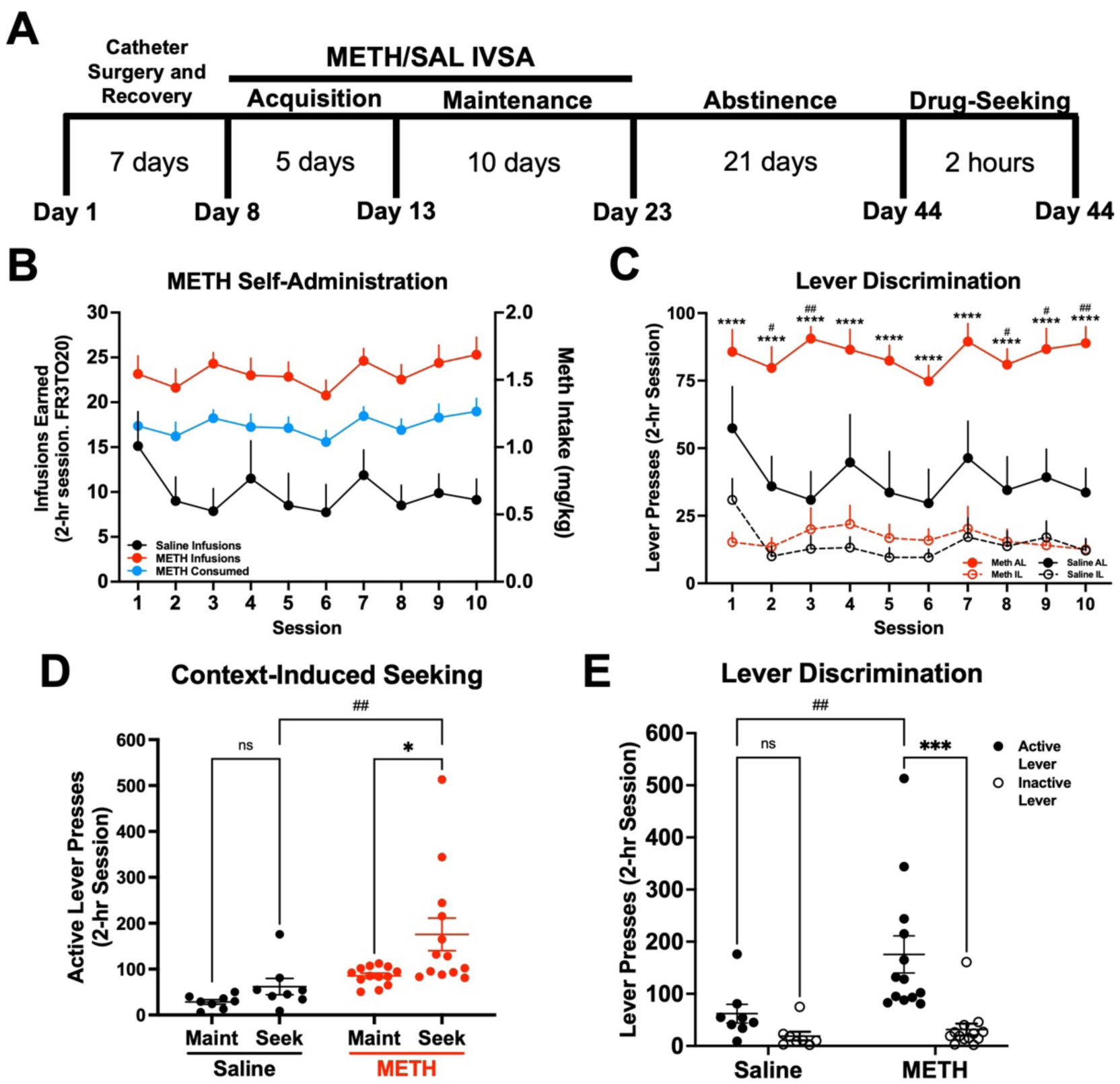
Establishment of a mouse model of METH IV self-administration. Male C57BL/6J mice were trained to self-administer METH (n = 13) or Saline (n = 8) during daily 2-hr sessions at FR3TO20. **A)** Experimental timeline. **B)** METH and Saline infusions earned, and METH consumed, during 15 daily 2-hr sessions (FR3TO20). **C)** Active and inactive lever presses during Maintenance. Two-way RM ANOVA with Bonferroni’s post-hoc test (Active vs Inactive Lever, \*\*\*\**p* < .0001; METH vs Saline Active Lever, ^#^*p* < .05, ^##^*p* < .01). **D)** Active lever presses for Maintenance (**Maint**: average final 3 days) and METH or Saline-seeking (**Seek**). Two-way RM ANOVA with Bonferroni post-hoc test (METH Maint vs Seek \**p* < .05; METH vs Saline Seek ^##^*p* < .01). **E)** Active and inactive lever presses during Drug-Seeking session. Two-way ANOVA with Bonferroni’s post-hoc test (METH Active vs Inactive Lever, \*\*\**p* < .001; METH Active vs Saline Active, ^##^*p* < .01). Data are represented as mean ± SEM.

### 3.2. Methamphetamine self-administration induces persistent transcriptional changes on dorsal striatal microglia

Microglia tightly regulate their gene expression in response to their environment (Ayata, Badimon et al. 2018, Masuda, Sankowski et al. 2019, Yeh and Ikezu 2019). With a working model of METH IVSA, we next investigated how microglia in the dorsal striatum, a brain region known for its role in methamphetamine-related behaviors (Chang, Alicata et al. 2007, Li, Rubio et al. 2015), alter their transcriptome in response to METH (or saline) IVSA and following 21 days of forced home cage abstinence (**Fig. 2A**). Importantly, affinity purification using Cd11b-positive selection yielded a highly pure population of microglia (**Supplemental Fig. 1**). Furthermore, RNA-sequencing of isolated dorsal striatal microglia revealed that numerous significant differentially expressed genes (DEGs) in response to methamphetamine (**Fig. 2B-D**). Methamphetamine administration induced more significantly upregulated than downregulated genes in dorsal striatal microglia (342 increased vs 190 decreased; Maintenance vs Saline, adjusted *p-value* < .05 and L2FC > 1.3 (**Fig. 2B**). Additionally, prolonged abstinence resulted in a similar number of significantly up- and downregulated genes (316 increased vs 358 decreased; Abstinence vs Maintenance, adjusted *p-value* < .05 and L2FC > 1.3 or L2FC < −1.3 (**Fig. 2C**). Notably, many genes following 21 days of abstinence were significantly upregulated than downregulated (240 increased vs 69 decreased; Abstinence vs Saline, adjusted *p-value* < .05 and L2FC > 1.3 (**Fig. 2D**). Gene expression across highly differentially expressed genes show similarity amongst samples from the same groups (**Fig. 2E**). Importantly, hierarchical clustering of samples within each condition based on DE genes indicates that mice exposed to methamphetamine (Maintenance and Abstinence) cluster more closely than to saline (**Supplementary Fig. 3**). These findings suggest methamphetamine significantly alters the transcriptome of dorsal striatal microglia. Considering that microglial function is directly tied to their gene expression and adapts to the shifting neural environment, we sought to identify the biological pathways related to the transcriptional differences in response to methamphetamine administration.

**Figure 2.**
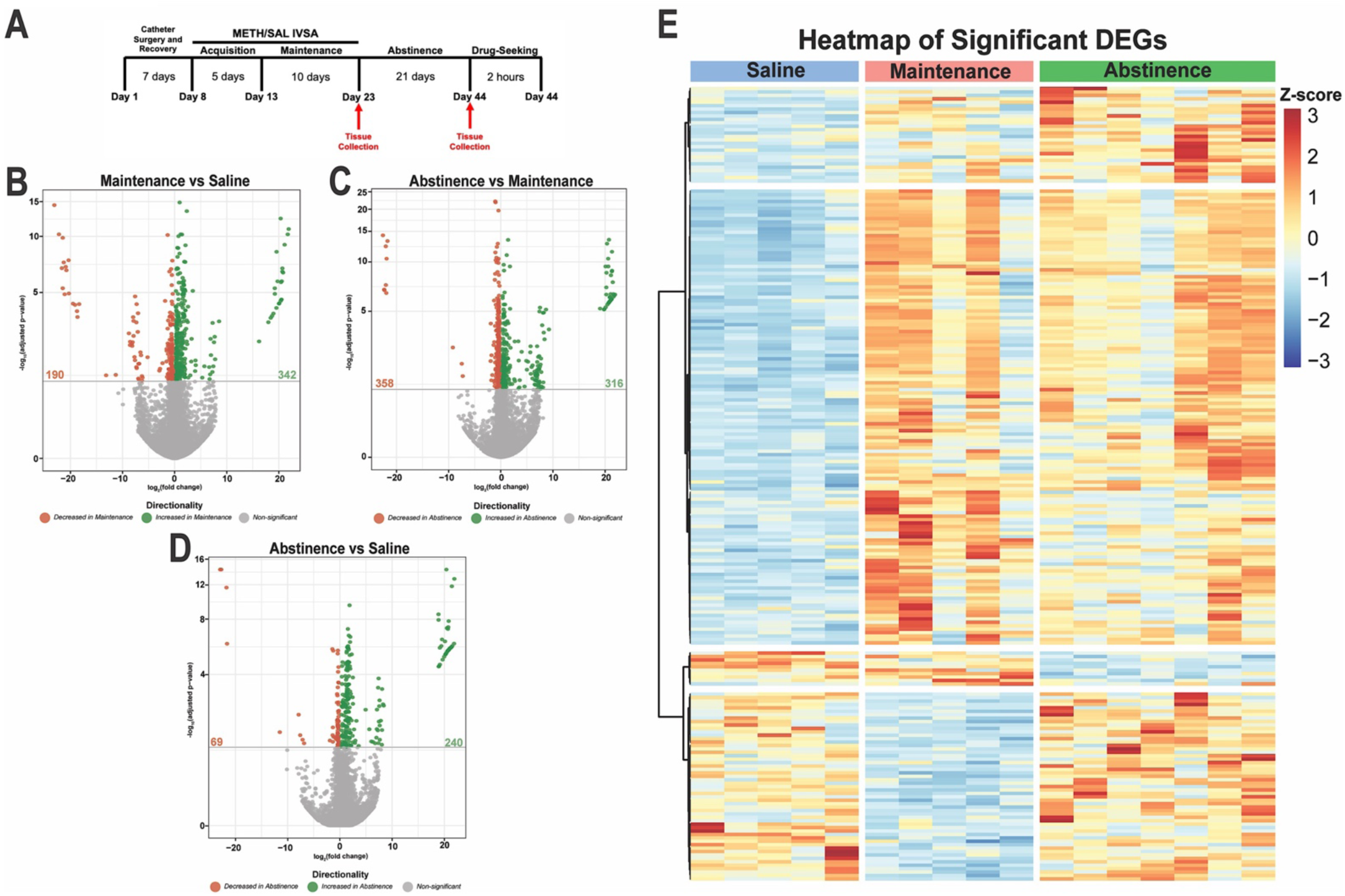
METH self-administration induces distinct transcriptional profiles in dorsal striatal microglia. **A)** Experimental timeline. Volcano plots showing significant DE genes (Log2FC vs −log_10_ adjusted *p-value*) in **B)** Maintenance vs Saline, **C)** Abstinence vs Maintenance, and **D)** Abstinence vs Saline. Red (decreased) and green (increased) circles represent DE genes that reached significance based on adjusted *p-value* < .05. **E)** Heatmap of normalized counts (rlog transformed) of selected significant DE genes (adjusted *p-value* < .05, lfcSE ≤ 1.5, L2FC ≤ −1.3 and L2FC ≥ 1.3) for each condition clustered by gene. Each column represents a single animal (Saline, blue; Maintenance, red; Abstinence, green). Saline (n = 5), Maintenance (n = 5), Abstinence (n = 7).

To this end, Gene ontology (GO) pathway analysis of significant DE genes revealed enrichment of biological pathways related to protein folding, mRNA processing, and cytoskeleton organization due to methamphetamine-taking (Maintenance vs Saline) (**Fig. 3A**). Of note, methamphetamine administration increased the expression of several heat shock proteins (e.g., Hspa8, Hspd1, Cryab, Ahsa2, and Dnaja1) (**Fig. 3B**). When comparing microglia from methamphetamine-abstinent to methamphetamine-taking mice (Abstinence vs Maintenance), pathways related to immune signaling and cellular stress response (e.g., apoptosis and response to radiation) were enriched (**Fig. 4A**). More specifically, compared to methamphetamine-taking mice, dorsal striatal microglia from methamphetamine-abstinent mice showed decreased expression of multiple heat shock proteins and apoptosis-related genes (e.g., Bax, Ddit3, Bclaf1, Acin1), yet increased expression of oxidative stress-related genes (e.g., Cirbp, Fus, and Kcnb1), and dysregulated immune signaling (e.g., Cd276, Nr1d1) (**Fig. 4B**). Additionally, pathways related to synapse organization and nervous system development were enriched in methamphetamine-abstinent mice (Abstinence vs Saline) (**Fig. 5A**). While heat shock proteins were generally downregulated when comparing Abstinence to Maintenance, several were still upregulated when comparing Abstinence to Saline (e.g., Cryab, Hspa1a, Dnaja1). Lastly, dorsal striatal microglia from methamphetamine-abstinent mice exhibited dysregulation of genes related to microglial activation (e.g., Syt11, Clu, Nr1d1) and upregulation of cell adhesion and morphology-related genes (e.g., Adora1, Ddr1, Tppp) (**Fig. 5B**). Overall, these findings suggest dorsal striatal microglia exhibit a unique transcriptome in response to methamphetamine administration and adopt a neuroprotective phenotype promoting neurogenesis, cell survival, and resolution of neuroinflammation after prolonged abstinence.

**Figure 3.**
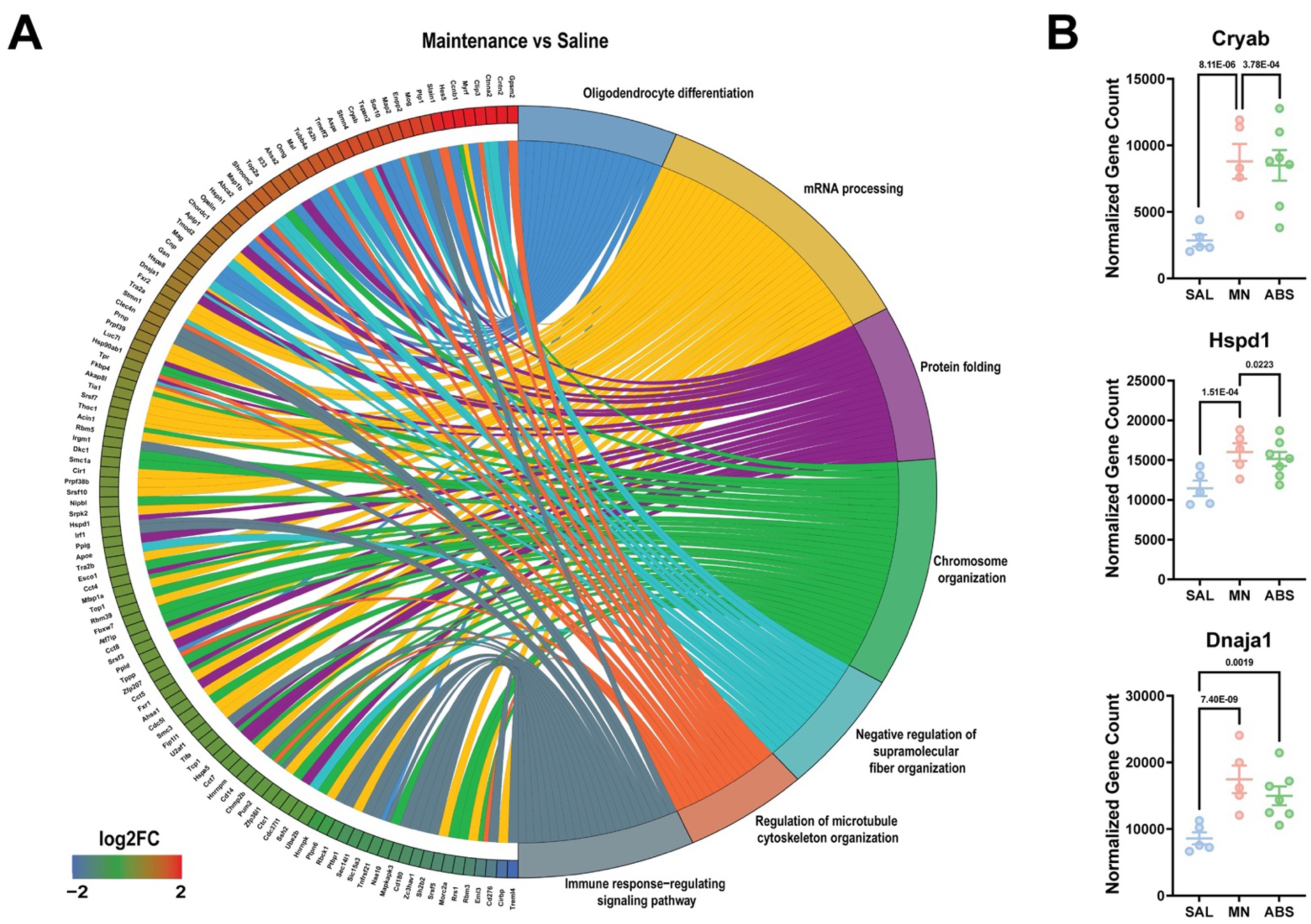
Gene Ontology (GO) enrichment analysis reveals dysregulation of protein folding and mRNA processing following METH self-administration. **A)** Chord plot showing interaction between GO biological process terms and genes comparing Maintenance vs Saline. **B)** Normalized counts of significant DE genes with adjusted *p-value* for each comparison. Significance shown reflects pairwise comparison results from DESeq2. Saline (n = 5), Maintenance (n = 5), Abstinence (n = 7). Data are represented as mean ± SEM.

**Figure 4.**
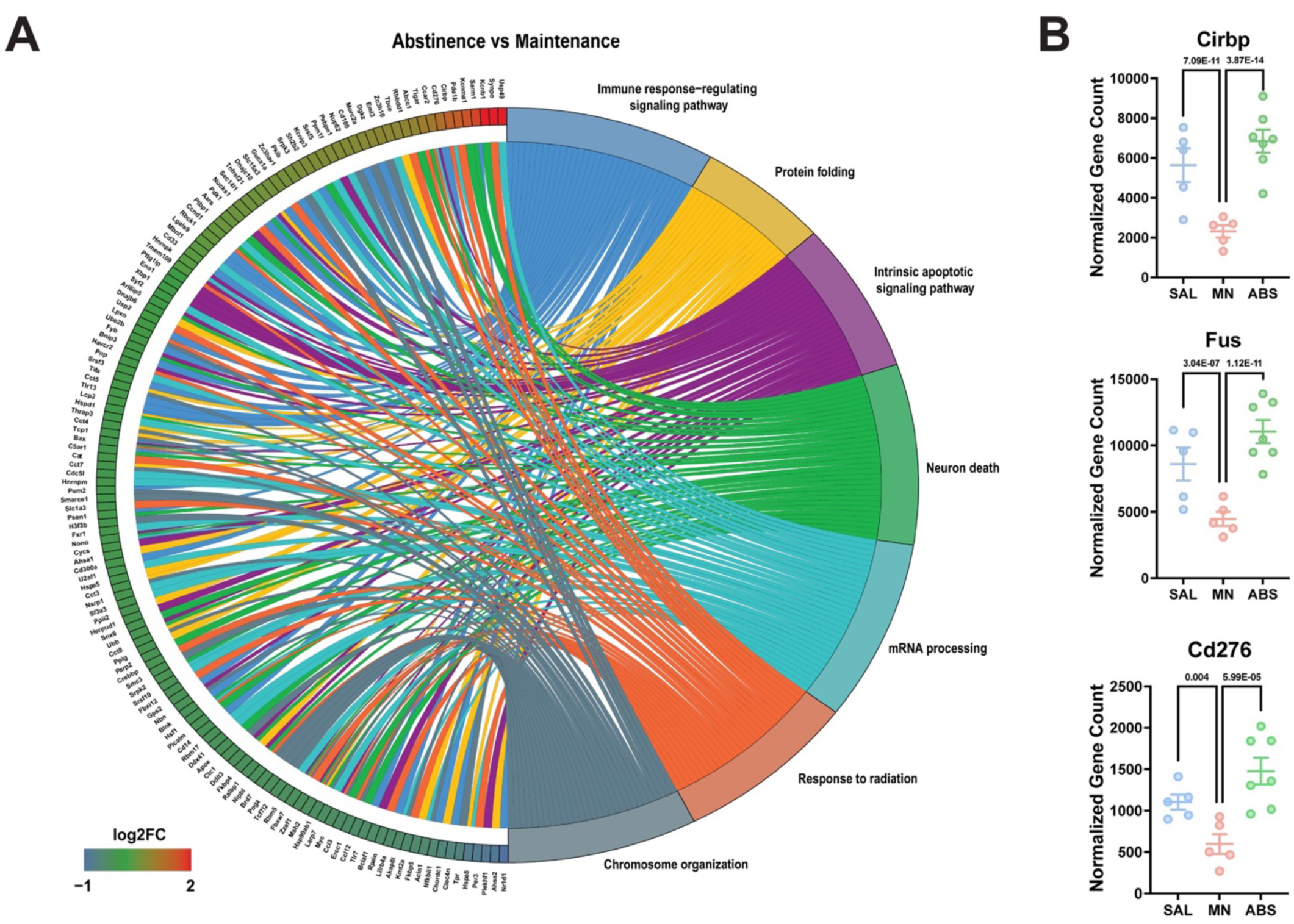
Gene Ontology (GO) enrichment analysis reveals dysregulation of immune signaling and cellular stress response in microglia previously exposed to METH. **A)** Chord plot showing interaction between GO biological process terms and genes comparing Abstinence vs Maintenance, **B)** Normalized counts of significant DE genes with adjusted *p-value* for each comparison. Significance shown reflects pairwise comparison results from DESeq2. Saline (n = 5), Maintenance (n = 5), Abstinence (n = 7). Data are represented as mean ± SEM.

**Figure 5.**
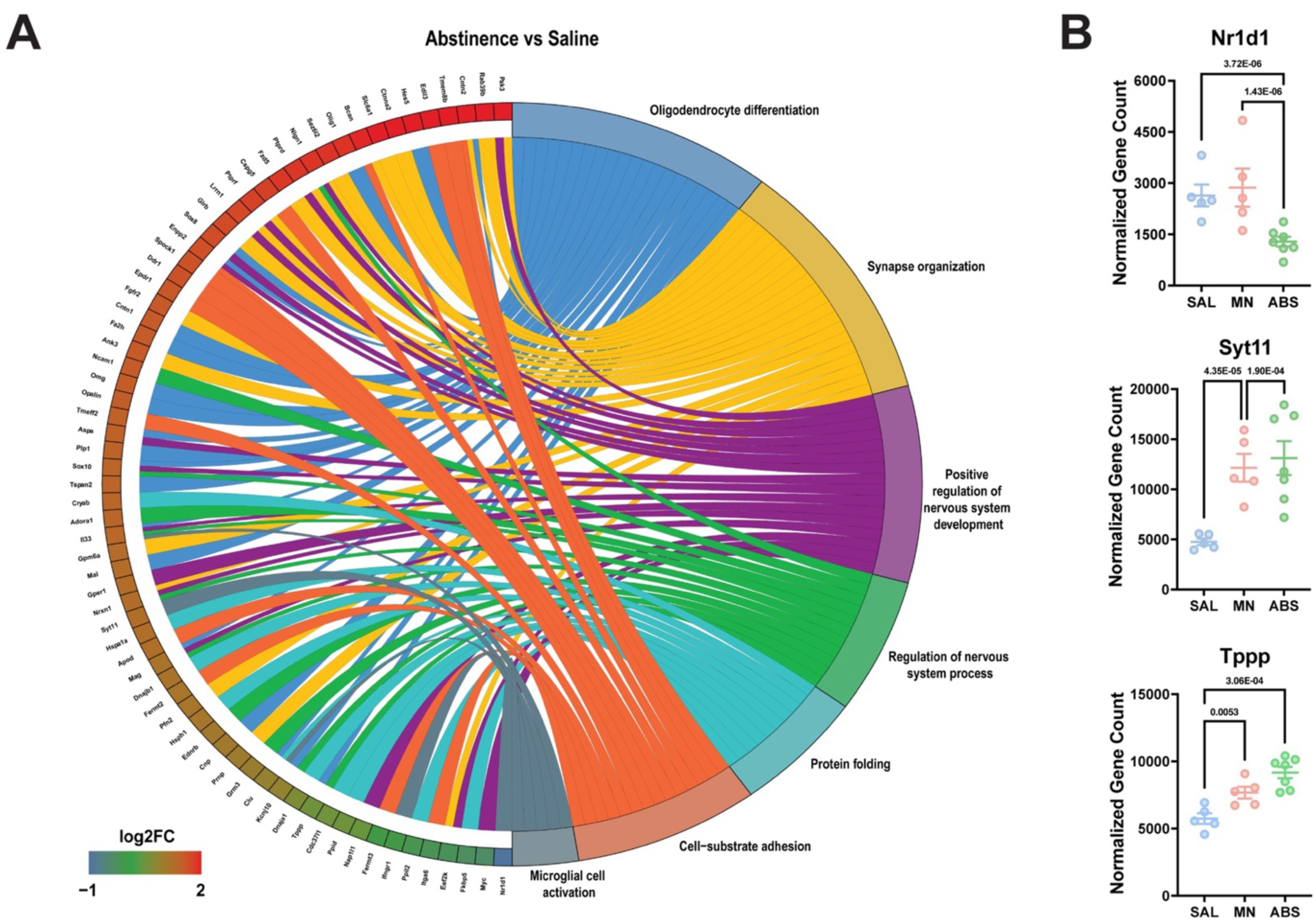
Gene Ontology (GO) enrichment analysis reveals persistent dysregulation of cell adhesion and microglial activation during abstinence. **A)** Chord plot showing interaction between GO biological process terms and genes comparing Abstinence vs Saline, **B)** Normalized counts of significant DE genes with adjusted *p-value* for each comparison. Significance shown reflects pairwise comparison results from DESeq2. Saline (n = 5), Maintenance (n = 5), Abstinence (n = 7). Data are represented as mean ± SEM.

### 3.3. Methamphetamine administration induces lasting morphological changes in dorsal striatal microglia

Microglial activity can be reflected in their morphology (Savage, Carrier et al. 2019, Vidal-Itriago, Radford et al. 2022). Specifically, “surveilling” microglia assume a more ramified morphology, while “effector” microglia deviate from this morphology (e.g., ameboid, hyper-ramified, etc.) (Morrison, Young et al. 2017, Savage, Carrier et al. 2019). Furthermore, several studies have shown exposure to stimulants such as cocaine and methamphetamine affect microglial morphology, even following extended withdrawal (LaVoie, Card et al. 2004, Sekine, Ouchi et al. 2008, Coelho-Santos, Goncalves et al. 2012, Reverte, Marchetti et al. 2024). Therefore, secondary to the transcriptome analysis in the previous section, we sought to determine if dorsal striatal microglia show concomitant changes in their morphology during METH (or saline) IVSA and following prolonged abstinence (**Fig. 6** and **Supplementary Fig. 4**). Representative fluorescence images of dorsal striatal microglia are shown in **Fig. 6A**. We employed Sholl analysis, a method of quantifying branching complexity (Sholl 1953), to assess morphological changes. Sholl analysis revealed microglia in methamphetamine-taking (Maintenance) and methamphetamine-abstinent (Abstinence) mice were significantly less ramified than Saline-taking (Saline) mice (**Fig. 6B**) (Two-way ANOVA; Distance x Condition Interaction, F (44, 2875)= 4.54, *p* < .0001). Furthermore, methamphetamine administration reduced the branching complexity of dorsal striatal microglia (**Fig. 6C**) (One-way ANOVA; Maintenance vs Saline, *p* < .0001), as indicated by significantly fewer branches (**Fig. 6D**) (One-way ANOVA; Maintenance vs Saline, *p* < .0001) and shorter processes (**Fig. 6E**) (One-way ANOVA; Maintenance vs Saline, p < .0001). Notably, microglial density in the dorsal striatum did not change across conditions (**Fig. 6F**). Interestingly, this effect persisted 21 days into abstinence (One-way ANOVA; Abstinence vs Saline Total Intersections, *p* < .0001; Abstinence vs Saline Mean Intersections, *p* < .0001; Abstinence vs Saline Max Branch Length, *p* = .0003). Taken together, these data demonstrate that dorsal striatal microglia adopt an altered morphology in response to methamphetamine, and that these changes last for several weeks following final methamphetamine exposure. Furthermore, given the neurotoxic properties of methamphetamine (Asanuma, Miyazaki et al. 2004, Jayanthi, Daiwile et al. 2021) these results are consistent with previous findings illustrating microglial reactivity following methamphetamine administration (Thomas, Walker et al. 2004, Robson, Turner et al. 2013, Goncalves, Leitao et al. 2017) and abstinence (Sekine, Ouchi et al. 2008, Yu, Chen et al. 2023).

**Figure 6.**
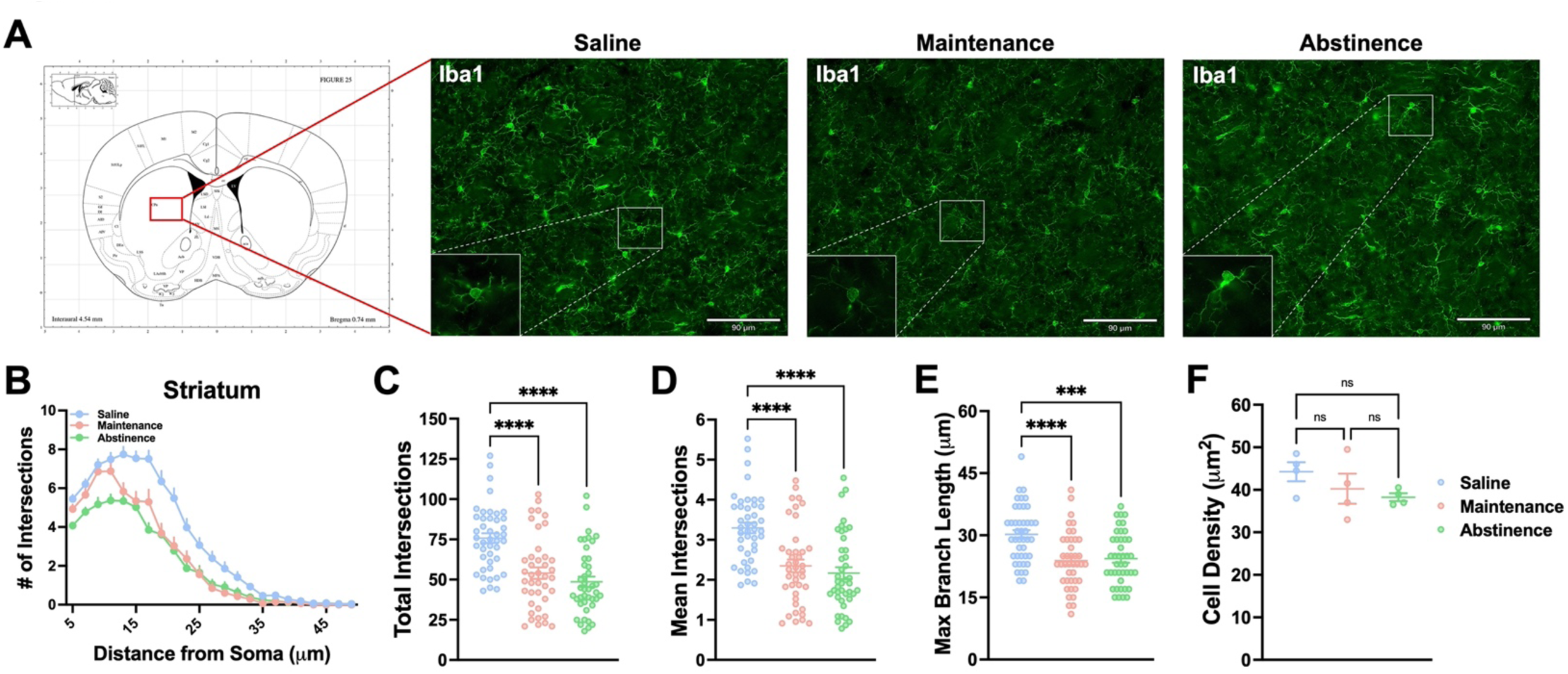
Dorsal striatal microglia exhibit persistent altered morphology following METH self- administration. **A)** Representative fluorescent images (Bregma 0.7-0.8 mm, Paxinos and Franklin’s the Mouse Brain in Stereotaxic Coordinates) of microglia (Iba1^+^ cells) in the dorsal striatum. **B)** Sholl analysis plot of microglia. **C-E)** Mice that self-administered METH (Maintenance), as well as METH-abstinent mice (Abstinence), display less ramifications and branching complexity than Saline-taking mice (Saline) with no significant change in density. One-way ANOVA with Tukey post-hoc test (Between Conditions, \*\**p* < .01, \*\*\**p* < 001, \*\*\*\**p* < .0001). 40-43 cells per condition (n = 4 animals, 10-12 cells per animal). Data are represented as mean ± SEM. Scale bar = 90 µm.

### 3.4. Sustained microglial depletion using PLX5622 during active methamphetamine-taking increases drug-intake

Methamphetamine administration induced persistent transcriptional and morphological changes in dorsal striatal microglia, suggesting a possible role in drug-taking behavior. To this end, we depleted microglia with PLX5622 during methamphetamine-taking (Acquisition and Maintenance). We found that mice self-administered methamphetamine regardless of treatment (**Fig. 7B**) (Two-way RM ANOVA; AIN-76A vs PLX5622, F (1, 17) = 1.88, *p* = .187). However, while mice treated with control AIN-76A chow showed stable intake across Maintenance, mice treated with PLX5622 chow gradually increased their methamphetamine intake (**Fig. 7C**) (t-test with Welch’s correction; AIN-76A vs PLX5622, *p* = .036). Additionally, while both treatment groups demonstrated robust discrimination between active and inactive levers over time (**Fig. 7D**) (Two-way RM ANOVA; Interaction between Session and Lever, F (42, 476) = 1.83, *p* = .002), mice lacking microglia exhibited significantly greater active lever pressing than control mice by the end of Maintenance (**Fig. 7E**) (Two-way ANOVA; Active vs Inactive Lever, F (1, 34) = 48.41, *p* < .0001; AIN-76A vs PLX5622, F (1, 34) = 4.11, *p* = .05). These data show that microglial ablation using PLX5622 increases methamphetamine self-administration, and as such, suggests that microglia may regulate the reinforcing properties of methamphetamine.

**Figure 7.**
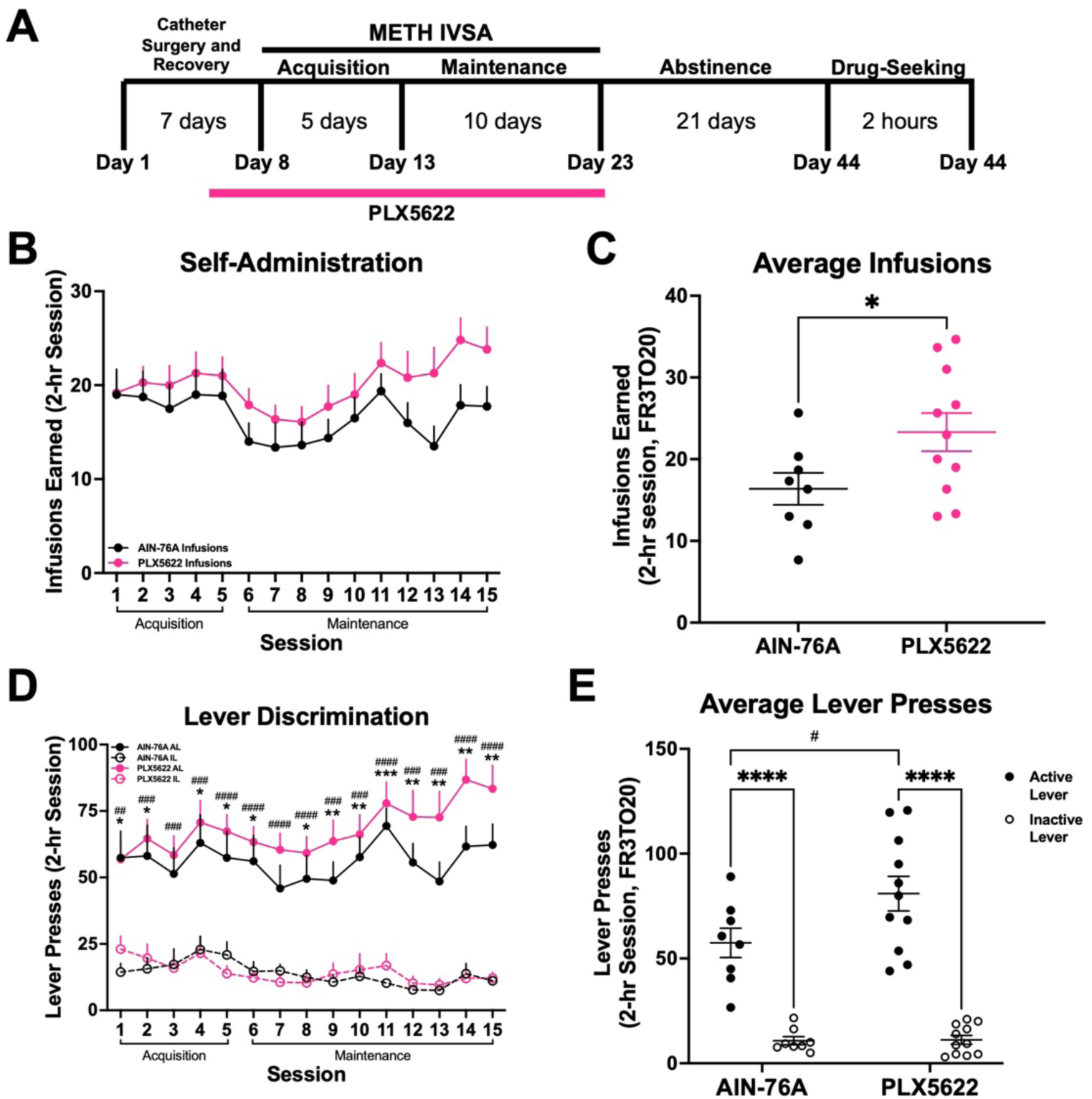
Microglial depletion increases METH self-administration. **A)** Experimental timeline. **B)** METH intake and infusions earned during 15 daily 2-hr sessions (FR3TO20). **C)** Average infusions earned over the last 3 days of Maintenance. Unpaired t-test with Welch’s correction (AIN-76A vs PLX5622, \**p* < .05). **D)** Active and inactive lever presses during 15 daily 2-hour sessions (FR3TO20). Two-way RM ANOVA with Bonferroni’s post-hoc test, (AIN-76A Active vs Inactive Lever, \**p* < .05, \*\**p* < .01, \*\*\**p* < .001; PLX5622 Active vs Inactive Lever, ^#^*p* < .05, ^##^*p* .01, ^###^*p* < .001, ^####^*p* < .0001) **E)** Average active and inactive lever presses over the last 3 days of Maintenance. Two-way ANOVA with Bonferroni’s post-hoc test (Active vs Inactive Lever, \*\*\*\**p* < .0001; AIN-76A Active vs PLX5622 Active, ^#^*p* < .05). AIN-76A (n = 8), PLX5622 (n = 11). Data are represented as mean ± SEM.

### 3.5. Treatment with PLX5622 during forced home cage abstinence does not affect context-induced methamphetamine-seeking

Since we found microglia to contribute to methamphetamine-taking behavior, we sought to determine if microglia contributed to methamphetamine-seeking. To this end, having completed 15 consecutive days of METH IVSA, mice were assigned to treatment groups (AIN-76A or PLX5622) for the duration of 21-day home cage abstinence (**Fig. 8A**). Importantly, treatment groups did not differ in number of methamphetamine infusions earned (**Fig. 8B**) (t-test with Welch’s correction; AIN-76A vs PLX5622, *p* = .572) or lever discrimination (**Fig. 8C**) (Two-way ANOVA; Active vs Inactive Lever, F (1, 32) = 109.3, *p* < .0001; AIN-76A vs PLX5622, F (1, 32) = .30, *p* = .587) prior to abstinence (combined data shown). We found that treatment with PLX5622 failed to significantly attenuate context-induced drug-seeking following forced home cage abstinence (**Fig. 8D**) (Two-way ANOVA; Maint vs Seek, F (1, 16) = 16.29, *p* = .001; AIN-76A vs PLX5622, F (1, 16) = .0617, *p* = .807). Additionally, both treatment groups displayed significant lever discrimination during the drug-seeking session (**Fig. 8E**) (Two-way ANOVA; Active vs Inactive Lever, F (1, 32) = 46.16, p < .0001; AIN-76A vs PLX5622, F (1, 32) = .2031, p = .861), suggesting a learned association between active lever pressing and drug-seeking. Thus, these results suggest that while microglia regulate the reinforcing properties of methamphetamine-taking (**Fig. 7B, C**), these cells may not be necessary for methamphetamine-seeking following prolonged abstinence.

**Figure 8.**
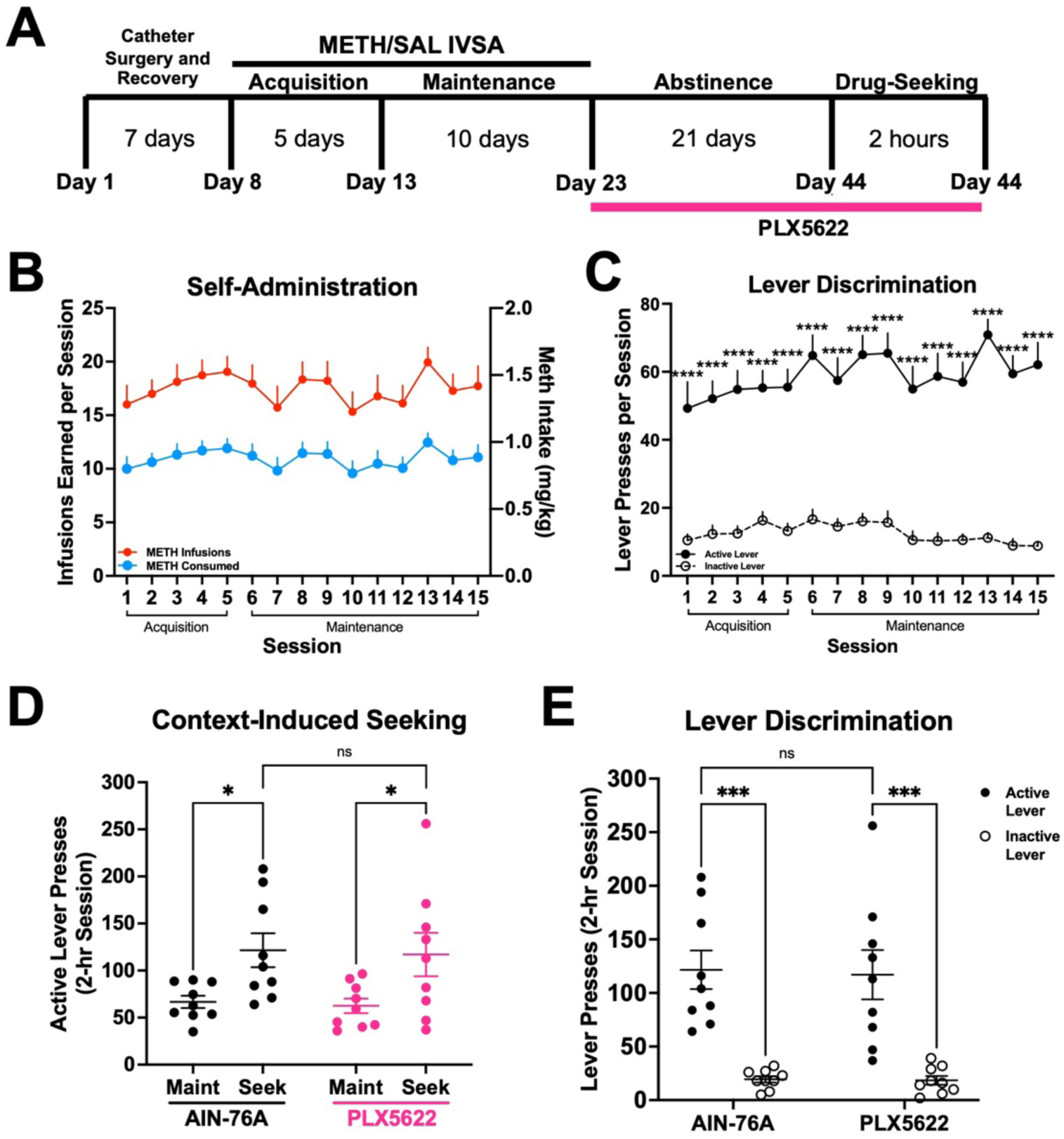
Microglial depletion during abstinence does not affect context-induced drug-seeking. **A**) Experimental timeline. **B)** METH intake and infusions earned during 15 daily 2-hr sessions. **C)** Active and inactive lever presses. Two-way RM ANOVA with Bonferroni’s post-hoc test, (Active vs Inactive Lever, \*\*\*\**p* < .0001). **D)** Active lever presses for Maintenance (**Maint**: average final 3 days) and Drug-Seeking (**Seek**) of AIN-76A and PLX5622. Two-way RM ANOVA with Bonferroni post-hoc test (AIN-76A Maint vs Seek, \**p* < .05; PLX5622 Maint vs Seek, \**p* < .05; AIN-76A vs PLX5622 Seek, *p* = .8). **E)** Active and inactive lever presses during Drug-Seeking session. Two-way ANOVA with Bonferroni’s post-hoc test (Active vs Inactive Lever, \*\*\**p* < .001, AIN-76A vs PLX5622 Active Lever*, p* = .853). AIN-76A (n = 9), PLX5622 (n = 9). Data are represented as mean ± SEM.

## 4. Discussion

### 4.1. Dorsal striatal microglia adopt a unique transcriptome and alter their morphology in response to methamphetamine administration and following prolonged abstinence

Consistent with the literature, we find that methamphetamine administration results in upregulation of gene expression associated with oxidative stress (Kuhn, Francescutti-Verbeem et al. 2006, Limanaqi, Gambardella et al. 2018, Yang, Wang et al. 2018). Specifically, methamphetamine administration resulted in a robust increase in heat shock protein expression, which has been linked to methamphetamine-induced hyperthermia (Cruickshank and Dyer 2009, Kiyatkin and Sharma 2011, Liao, Lu et al. 2021) and the production of reactive oxygen species from methamphetamine-induced neurotoxicity and terminal degeneration in the striatum (Asanuma, Miyazaki et al. 2004, McConnell, O’Banion et al. 2015, Frank, Adhikary et al. 2016). Importantly, many of these genes remained dysregulated following 21 days of abstinence, which is consistent with human PET studies where microglial activation persists 2 years after methamphetamine cessation (Sekine, Ouchi et al. 2008). Consistent with upregulation of genes related to cell adhesion (e.g., Tppp, Adora1, Tubb4a), morphological analysis revealed that dorsal striatal microglia have reduced branching complexity that remains through abstinence. Additionally, we found that following methamphetamine administration, dorsal striatal microglia share similar gene expression to disease-associated microglia from neurodegenerative diseases such as Alzheimer’s (Corneveaux, Myers et al. 2010) and Parkinson’s disease (PD) (Du, Wang et al. 2017). Notably, the upregulation of several genes associated with PD (e.g., Cryab, Syt11, Hspa8, Stip1) in our dataset support studies indicating individuals with methamphetamine use disorder are three times more likely to develop PD (Callaghan, Cunningham et al. 2012, Curtin, Fleckenstein et al. 2015), and suggest microglia may contribute to this increased risk. These results further suggest methamphetamine administration induces lasting effects on the neuronal environment and that microglia adapt to these environmental changes by altering their transcriptome, which is reflected in their morphology.

Consistent with our findings, other studies indicate that psychomotor stimulants such as methamphetamine and cocaine persistently alter microglial morphology (LaVoie, Card et al. 2004, Sekine, Ouchi et al. 2008, Coelho-Santos, Goncalves et al. 2012, Reverte, Marchetti et al. 2024). In fact, recent evidence suggests that microglia adopt a wide array of morphologies in response to various stimuli (Fontainhas, Wang et al. 2011, Morrison, Young et al. 2017, Vidal-Itriago, Radford et al. 2022), often exhibiting both hyper-ramification and de-ramification within the same region (Morrison and Filosa 2013). Therefore, further studies are required to characterize the morphological heterogeneity of microglia in response to methamphetamine administration. By complementing morphological analyses with cell-specific RNA-sequencing, our findings demonstrate that microglia may adopt a more anti-inflammatory (neuroprotective) state during abstinence by modulating their gene expression in favor of nervous system development and repair (e.g., Ptgds, Nrxn1, Cntn1, Lgals1) (Colton 2009, Starossom, Mascanfroni et al. 2012, Hickman, Kingery et al. 2013), as well as promote their own survival and proliferation (e.g., Dnaja1, Bnip3, Acin1, Hspa1a, Clu), indicating that microglia may assist in the resolution of inflammation and restoration of homeostasis in the neural environment of the dorsal striatum.

Considering the persistence of these transcriptional and morphological changes, our data suggest that epigenetic mechanisms may also be involved in their perpetuation. Although we did not find numerous epigenetic gene-related expression changes within our dataset, several studies have highlighted the importance of epigenetic regulation of gene expression following methamphetamine administration (Omonijo, Wongprayoon et al. 2014, Cadet, Brannock et al. 2015) and microglial activity (Matcovitch-Natan, Winter et al. 2016, Ayata, Badimon et al. 2018, Cheray and Joseph 2018). Specifically, H3K4 methylation and H3K27 acetylation, markers of active gene promoters and enhancers (Calo and Wysocka 2013), are linked to innate immune memory in macrophages (Kaikkonen, Spann et al. 2013, Ostuni, Piccolo et al. 2013), and more recently in microglia (Meleady, Towriss et al. 2023). These histone post-translational modifications, along with other epigenetic enzymes and chromatin remodeling complexes, have implicated microglia in the progression of neurodegenerative disease and aging (Cho, Chen et al. 2015, Yeh and Ikezu 2019, Huang, Malovic et al. 2023), as well as substance use disorders (Schwarz, Hutchinson et al. 2011, Crews, Coleman et al. 2023, Vilca, Margetts et al. 2023). Therefore, future studies will be needed to examine the underlying epigenetic machinery in microglia that may govern the lasting transcriptional and morphological changes effected by methamphetamine administration and during abstinence to the drug.

### 4.2. Are microglia protective against excessive methamphetamine-taking?

We found that methamphetamine administration is influenced by the absence of microglia, as treatment with PLX5622 during active methamphetamine-taking increased intake of the drug. Importantly, and consistent with other studies, the absence of microglia did not affect the cellular morphology of other cells within the dorsal striatum (**Supplementary Fig. 5**) (Zhan, Krabbe et al. 2019, Du, Brennan et al. 2022), nor did it affect operant responding for a natural reward (**Supplementary Fig. 6**), suggesting that microglia specifically regulate the reinforcing properties of methamphetamine in our behavioral model. Interestingly, we found several genes involved in neurotransmitter signaling and synthesis to be upregulated in response to methamphetamine administration and following abstinence (**Supplementary Figs. 7 and 8**), underscoring the neurotransmitter-sensing capabilities of microglia and their ability to modulate neuronal activity and neurotransmitter release (Badimon, Strasburger et al. 2020, Stolero and Frenkel 2021). For example, microglia are thought to contribute to the reuptake of GABA (Bhandage and Barragan 2021, Favuzzi, Huang et al. 2021), a neurotransmitter that is elevated in the striatum during administration of and withdrawal from psychostimulants (Wydra, Golembiowska et al. 2013). Indeed, we found the GABA transporter 1 (GAT1) to be significantly upregulated during methamphetamine-taking and abstinence (**Supplementary Fig. 7**). Furthermore, several genes related to glutamate synaptic clearance (GLT-1), signaling (mGluR3, Gria2, Glrb), and processing (Glul, Glud1, Got1, Gls) were also increased following methamphetamine administration and abstinence (**Supplementary Fig. 7**) (van Landeghem, Stover et al. 2001). As such, eliminating microglia may disrupt homeostatic responses to methamphetamine aimed at maintaining excitatory/inhibitory balance in the dorsal striatum. Additionally, methamphetamine-induced upregulation of dopamine signaling genes (Gpr37, Ddc, Darpp-32, Cdh11) in dorsal striatal microglia (**Supplementary Fig. 8**) further suggests that these cells may alter their gene expression to adapt to increased striatal dopamine normally seen following methamphetamine administration (Mark, Soghomonian et al. 2004). Therefore, depleting microglia may impair endogenous mechanisms to maintain neurotransmitter balance in the dorsal striatum – an effect that becomes evident as animals gradually escalate intake over repeated methamphetamine self-administration sessions (**Fig. 7B**). It should be noted that while the overall effect of microglial depletion could reflect increased motivation for the drug, it is also possible that this effect may be due to decreased sensitivity to the effects of methamphetamine. Further studies employing a progressive ratio/breakpoint paradigm (Caprioli, Zeric et al. 2015) and/or a dose-response curve (Munzar, Laufert et al. 1999, Calabrese and Baldwin 2001) will be able to determine the nature of increased methamphetamine-taking in the absence of microglia.

While PLX5622 treatment increased methamphetamine-taking, it was not sufficient to attenuate context-induced methamphetamine-seeking, suggesting that microglia may not play a determining role in methamphetamine-seeking when these cells are depleted during drug abstinence. This finding is consistent with literature demonstrating that chronic delivery of the TLR4 antagonist (+)-naltrexone does not affect cue-induced reinstatement of methamphetamine-seeking after 13 days of abstinence (Theberge, Li et al. 2013). Additionally, a separate study showed global knockout of TNF-α increased methamphetamine self-administration and motivation but did not affect cue-induced reinstatement (Yan, Nitta et al. 2012). Consistent with these findings, a recent report has shown that PLX5622-mediated microglial depletion during cocaine withdrawal can reduce conditioned hyperlocomotion without affecting drug memory (Reverte, Marchetti et al. 2024). However, while PLX5622 treatment during 21-day abstinence did not affect drug-seeking in our animals, we did observe that repopulation of microglia during abstinence prevented drug-seeking (**Supplementary Fig. 9**), indicating that indeed, microglia may play a role in methamphetamine-related drug memory. This finding is in line with other studies which have demonstrated that microglia are involved in memory formation and that microglial depletion using PLX5622 reduces dendritic spine development and alters behavior in memory-related tasks (Parkhurst, Yang et al. 2013, Basilico, Ferrucci et al. 2022, Reverte, Marchetti et al. 2024). Taken together, these data suggest microglia contribute to methamphetamine-induced neural adaptions, and that their role in methamphetamine reinforcement and seeking may be closely associated to the timepoint in the behavioral course of the disease.

### 4.3. Limitations of the current study

Sex is an important consideration when studying MUD (McHugh, Votaw et al. 2018). Women have been reported to use methamphetamine earlier in life and become more dependent (Dluzen and Liu 2008). Additionally, neuroimmune system development is regulated by sex (McCarthy, Nugent et al. 2017, Osborne, Turano et al. 2018). Consequently, a limitation of the current study was only using male mice for the behavioral and molecular experiments. Also of note, while the current study examined dorsal striatal microglia, as this brain region has been heavily implicated in the development and maintenance of MUD (Chang, Alicata et al. 2007), other brain regions such as the hippocampus (Goncalves, Baptista et al. 2010, Takashima, Fannon et al. 2018) and various cortical regions (Gonzalez, Jayanthi et al. 2018, Kearns, Siemsen et al. 2022) also contribute to methamphetamine-related behaviors and pathophysiology of MUD, as well as transcriptional differences in microglia (Barko, Shelton et al. 2022). Therefore, further studies focusing on sex differences and relevant brain regions beyond the dorsal striatum will be necessary to gain a better understanding of the underlying microglial mechanisms regulating methamphetamine reinforcement.

## 5. Conclusion

Our data suggest that microglia in the dorsal striatum adopt a persistent neuroprotective phenotype in response to methamphetamine administration. In addition, methamphetamine-induced dysregulation of GABA and glutamate neurotransmission genes suggest that microglia may also play a role in methamphetamine reinforcement beyond neuroimmune regulation, an effect supported by increased methamphetamine-taking in the absence of microglia. Altogether, this study increases our understanding of how microglia adapt their gene expression and morphology to methamphetamine administration and seeking and may provide insights into the role of microglia in the methamphetamine reinforcement and methamphetamine use disorder pathophysiology.

## Acknowledgments

SJV, AVM, and LMT designed and coordinated the study. SJV conducted all behavioral experiments and obtained samples for RNA sequencing and IHC. SJV and AVM processed samples for RNA sequencing, and SJV and IF processed samples for IHC. SJV and AVM conducted statistical analysis and data interpretation. SJV, AVM, and LMT drafted the manuscript. IF was supported by NCI grant R25CA261632. This work was supported by NIDA grants K01DA045294 (LMT), DP1DA051828 (LMT), U18DA052533 (CW), NINDS grant F99NS130871 (SJV), the NIDA Drug Supply Program, as well as a kind gift from the Shipley Foundation (LMT).

## Supplementary Methods and Results

### RNA-sequencing analysis of isolated dorsal striatal microglia

Biological replicates determined to be outliers were removed for differential gene expression analysis (**Supplementary Fig. 3A**). Principal component analysis (PCA) (**Supplementary Fig. 3B**) and heatmap of hierarchical clustering of conditions based on gene expression (**Supplementary Fig. 3C**) shows high similarity of samples within condition, and that animals exposed to methamphetamine (Maintenance and Abstinence) cluster more closely than to Saline.

### Microglia are not required for natural food reinforcement

To test if microglia are necessary for learned operant behavior, we food-trained mice up to FR5 for 8 consecutive days (**Supplementary Fig. 5**). Mice were treated with PLX5622 (1200 ppm in AIN-76A chow) for the duration of the experiment. Microglial ablation does not affect natural food reinforcement in number of rewards earned (**Supplementary Fig. 5A**) (Two-way RM ANOVA; AIN-76A vs PLX5622, F (1, 13) = .073, *p* = .791) or lever discrimination (**Supplementary Fig. 5B**) (Two-way RM ANOVA; Active vs Inactive Lever, F (3, 26) = 24.38, *p* < .0001) and time to acquire operant lever pressing behavior (**Supplementary Fig. 5B**) (Two-way RM ANOVA; AIN-76A vs PLX5622, F (1, 13) = .385, *p* = .545).

**Supplementary Figure 1.**
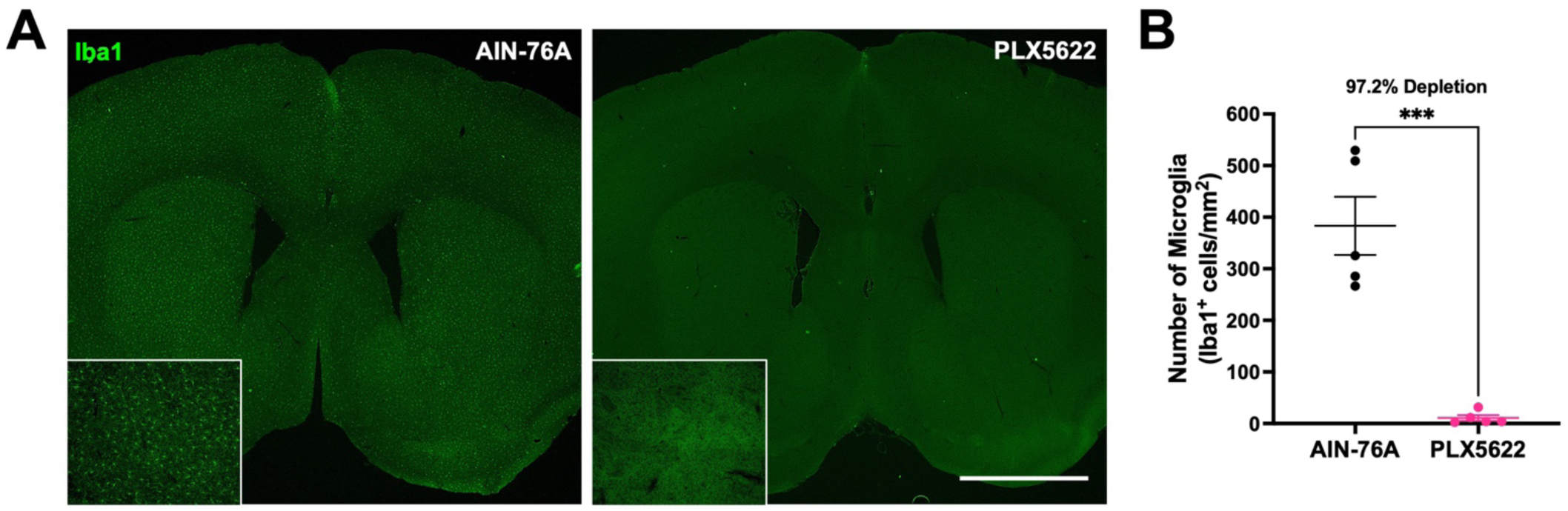
Treatment with CSF1R inhibitor PLX5622 results in near complete depletion of microglia. **A)** Representative fluorescent images of Iba1^+^ microglia (green) in the dorsal striatum from AIN-76A and PLX5622-treated mice. **B)** Quantification of microglial density. Unpaired t-test (AIN-76A vs PLX5622, \*\*\**p* < .001). n = 5 per group. Data are represented as mean ± SEM. Scale bar = 1360 μm.

**Supplementary Figure 2.**
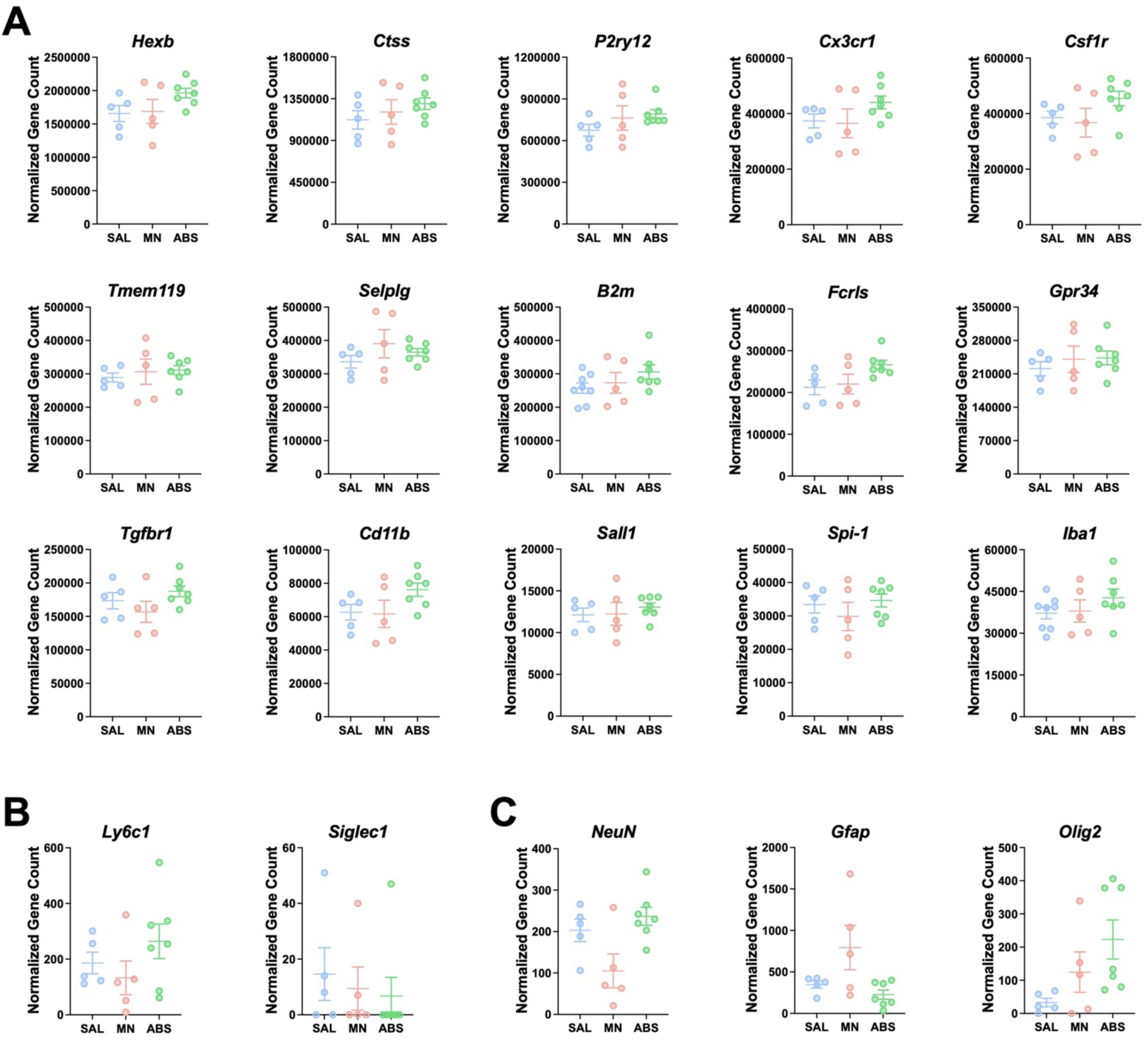
Purity of isolated dorsal striatal microglia. **A)** Normalized counts of microglia-specific genes. **B)** Normalized counts of macrophage-specific genes. **C)** Normalized counts of other neural cell types-specific genes: neurons (*NeuN*), astrocytes (*Gfap*), oligodendrocytes (*Olig2*).

**Supplementary Figure 3.**
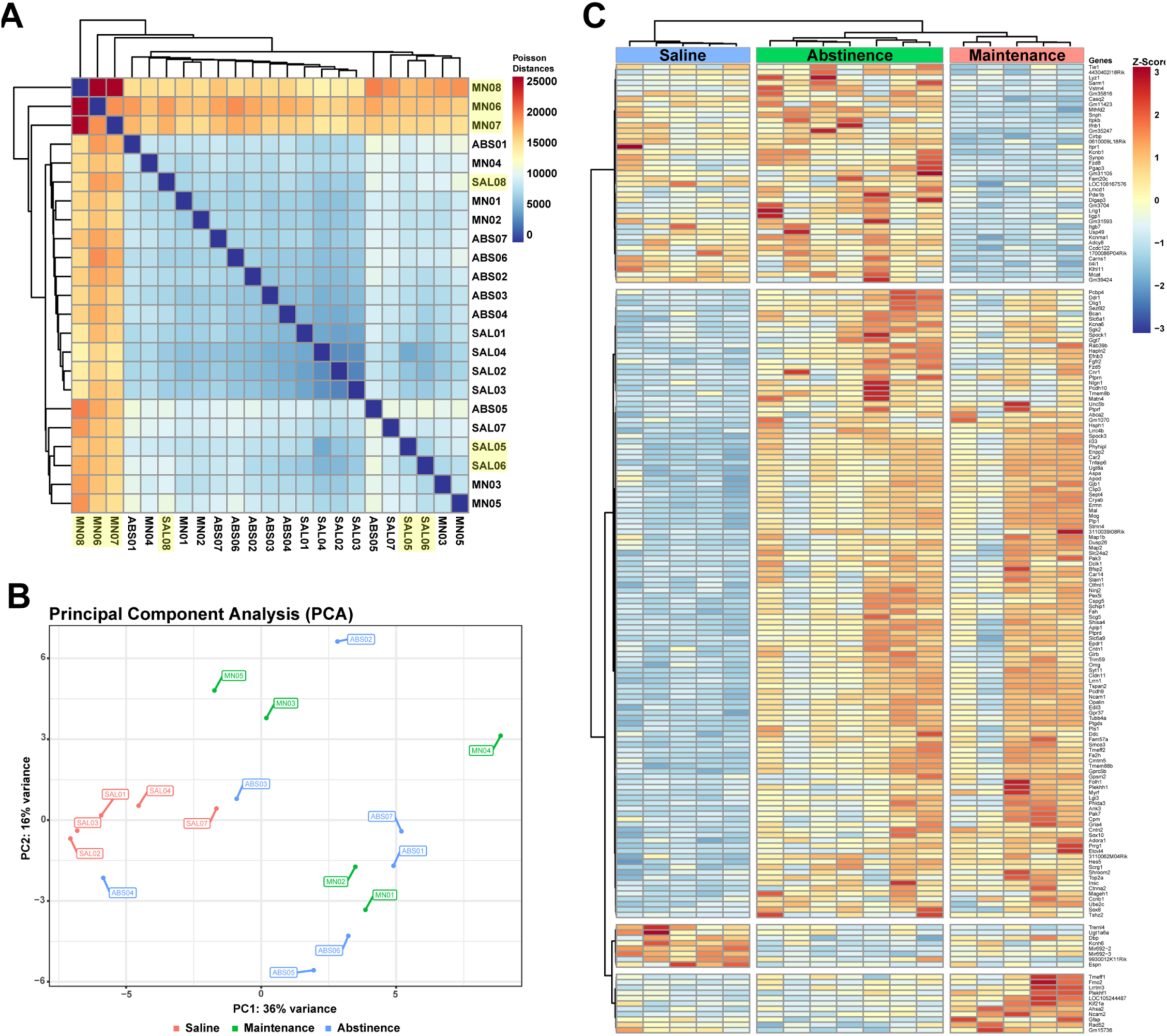
RNA-sequencing of isolated dorsal striatal microglia from METH IVSA. **A)** Hierarchical clustering heatmap of expression profiles for samples (n = 23) based on Poisson distance. Highlighted samples were determined to be outliers and were removed from analyses. **B)** PCA plot for samples (n = 17) following removal of outliers. **C)** Heatmap showing unsupervised clustering of samples based on gene expression.

**Supplementary Figure 4.**
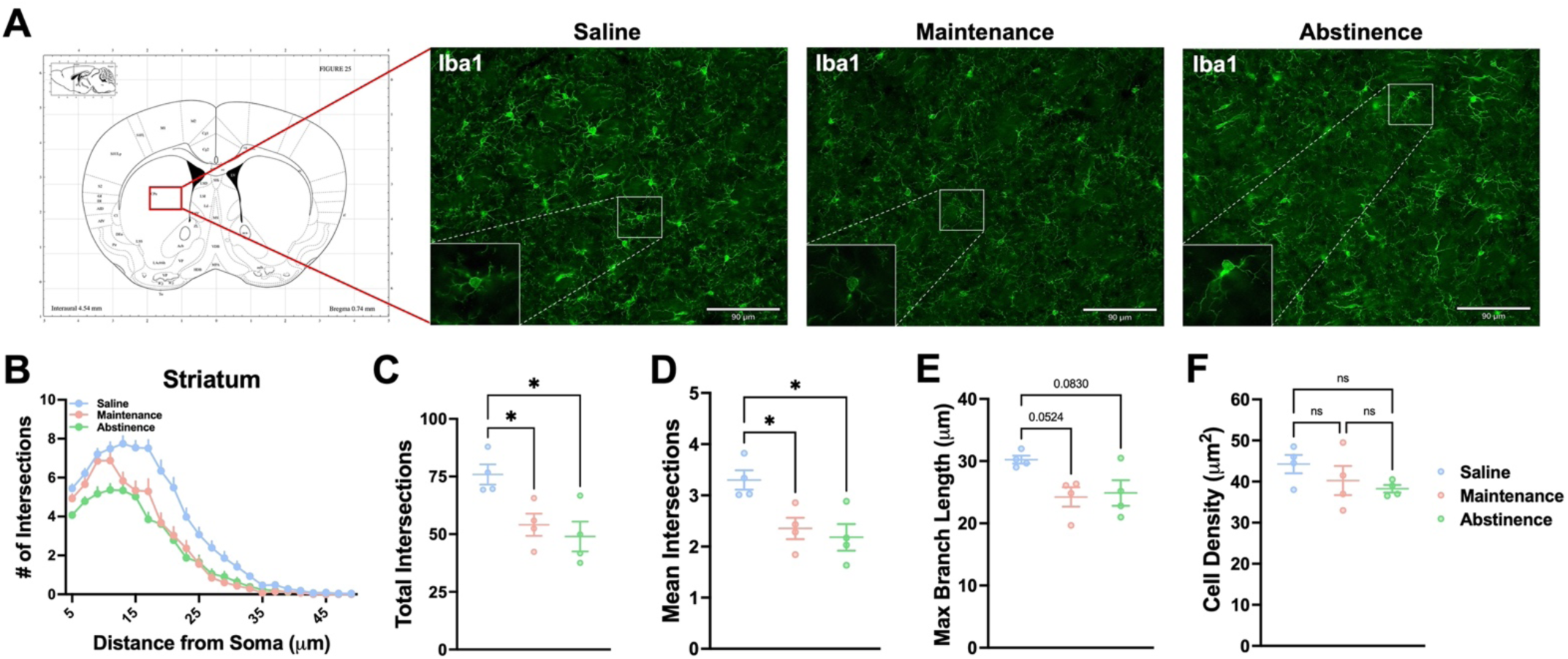
Dorsal striatal microglia show persistent altered morphology due to METH administration. **A)** Representative fluorescent images (Bregma 0.74 mm, Paxinos and Franklin’s the Mouse Brain in Stereotaxic Coordinates) of microglia (Iba1^+^ cells) in the striatum. **B)** Sholl analysis plot of microglia. **C-F)** Mice that self-administered METH (Maintenance), as well as METH-abstinent mice (Abstinence), display less ramifications and branching complexity and shorter processes than Saline-taking mice (Saline), with no significant change in density. One-way ANOVA with Tukey post-hoc test (between conditions, \**p* < .05). n = 4 animals for all conditions. Data are represented as mean ± SEM. Scale bar = 90 µm.

**Supplementary Figure 5.**
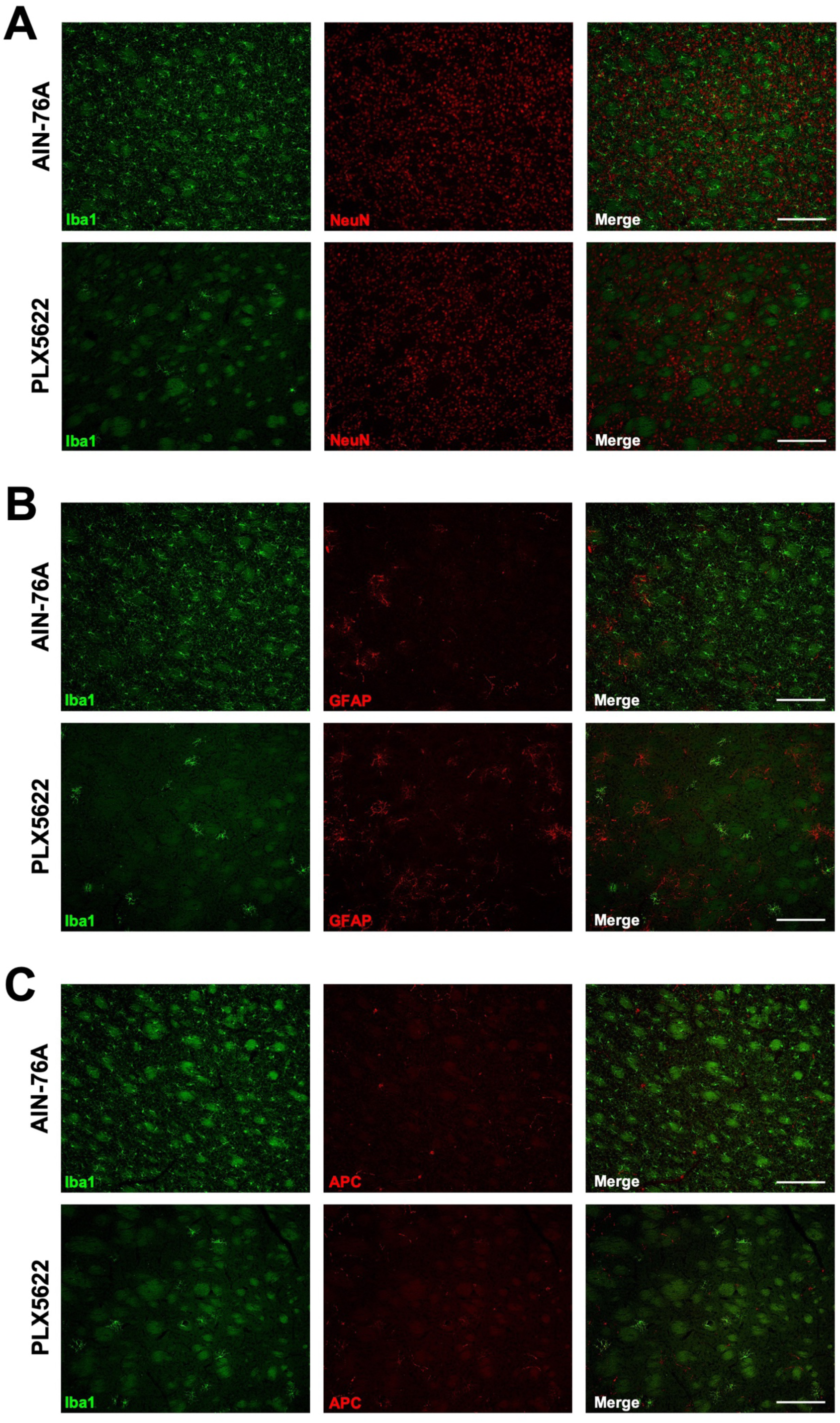
PLX5622 treatment does not affect general morphology of neural cells in the dorsal striatum. **A)** Representative 10X images of microglia (Iba1^+^, green) and neurons (NeuN^+^, red). **B)** Representative 10x images of microglia (Iba1^+^, green) and astrocytes (GFAP^+^, red). **C)** Representative 10X images of microglia (Iba1^+^, green) and oligodendrocytes (APC^+^, red). Scale bar = 170 μm.

**Supplementary Figure 6.**
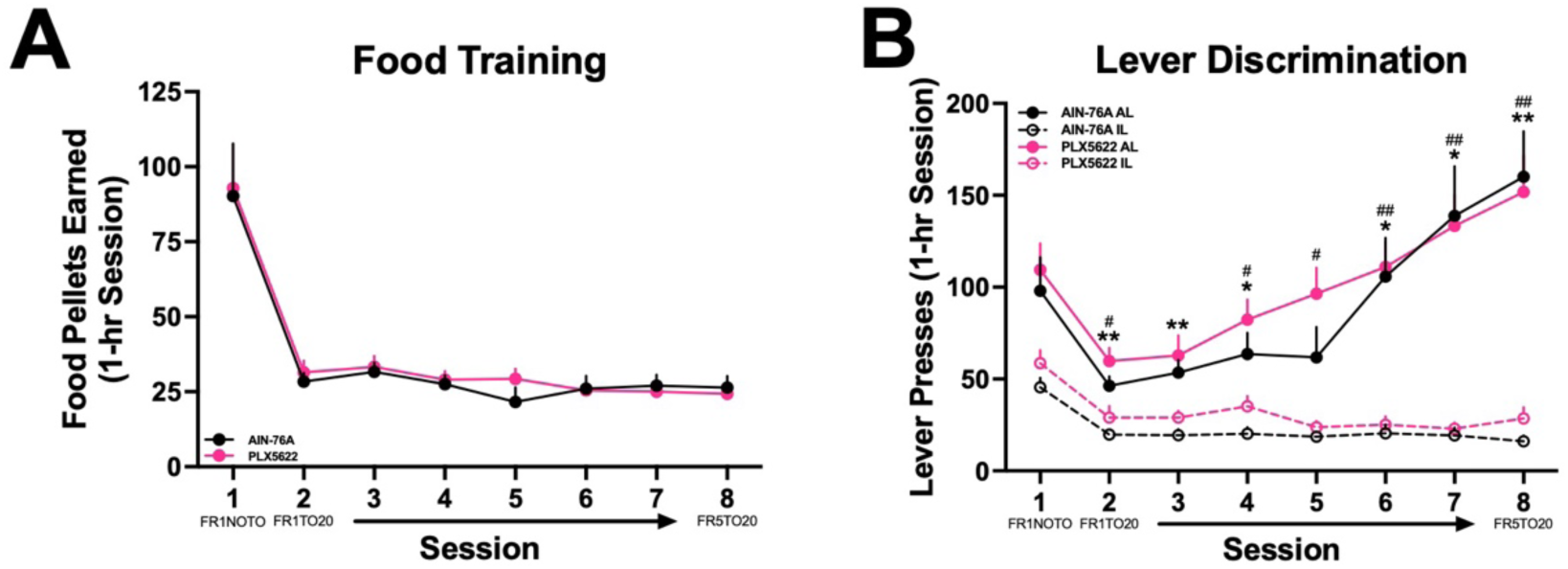
Pharmacological ablation of microglia does not affect operant responding. **A)** Number of food rewards earned during 8 daily 1-hr sessions. **(B)** Active vs inactive lever presses during 8 daily 1-hr sessions (Two-way RM ANOVA with Bonferroni post-hoc test; AIN-76A Active vs Inactive Lever, \**p* < .05, \*\**p* < .01; PLX5622 Active vs Inactive Lever, ^#^*p* < .05, ^##^*p* < .01). AIN-76A (n = 8), PLX5622 (n = 7). Data are represented as mean ± SEM.

**Supplementary Figure 7.**
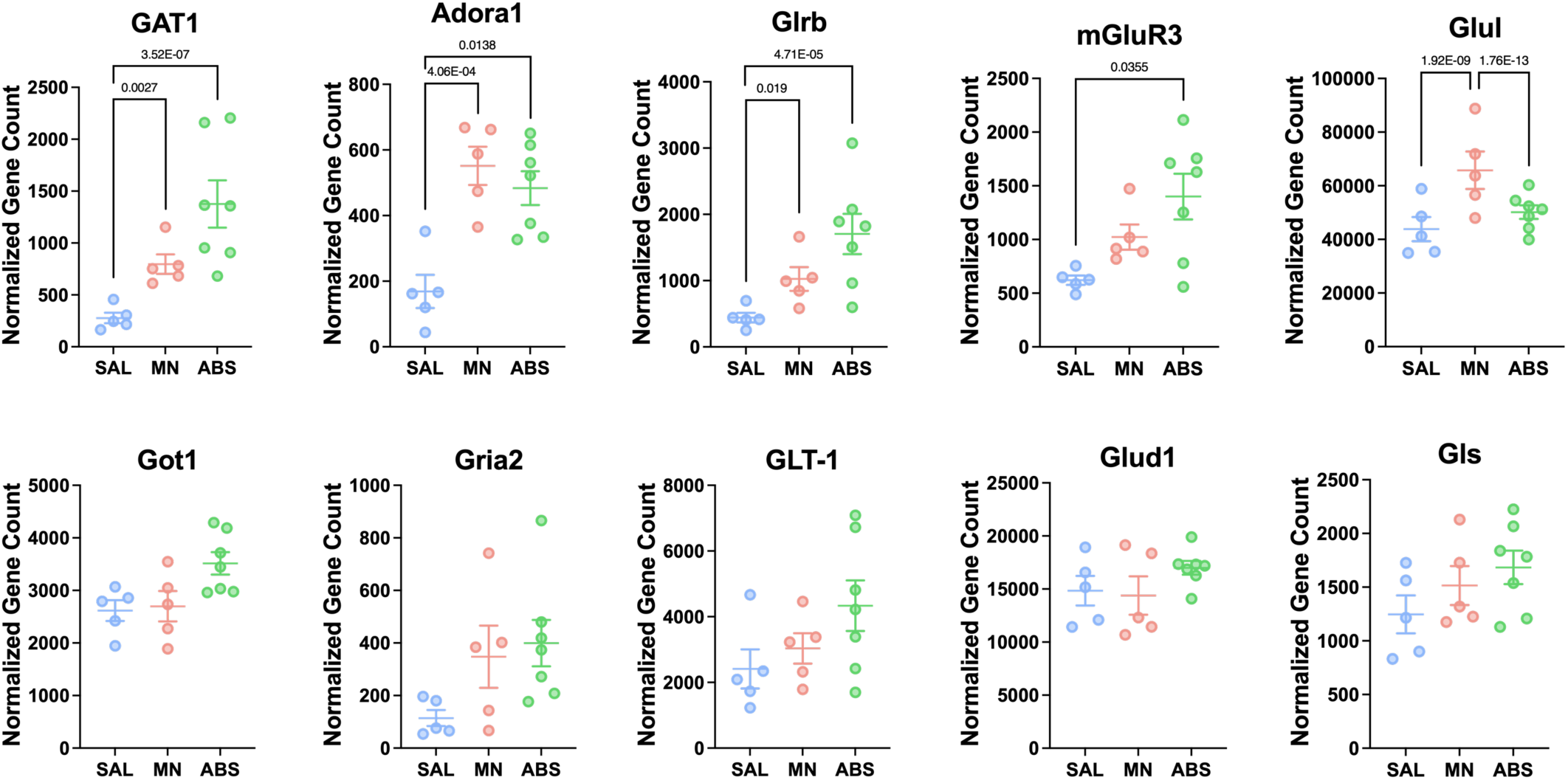
GABA, glutamate, and adenosine signaling-related genes. Normalized counts of DE genes related to GABA, glutamate, and adenosine signaling with adjusted *p-value* for each comparison. Significance shown reflects pairwise comparison results from DESeq2.

**Supplementary Figure 8.**
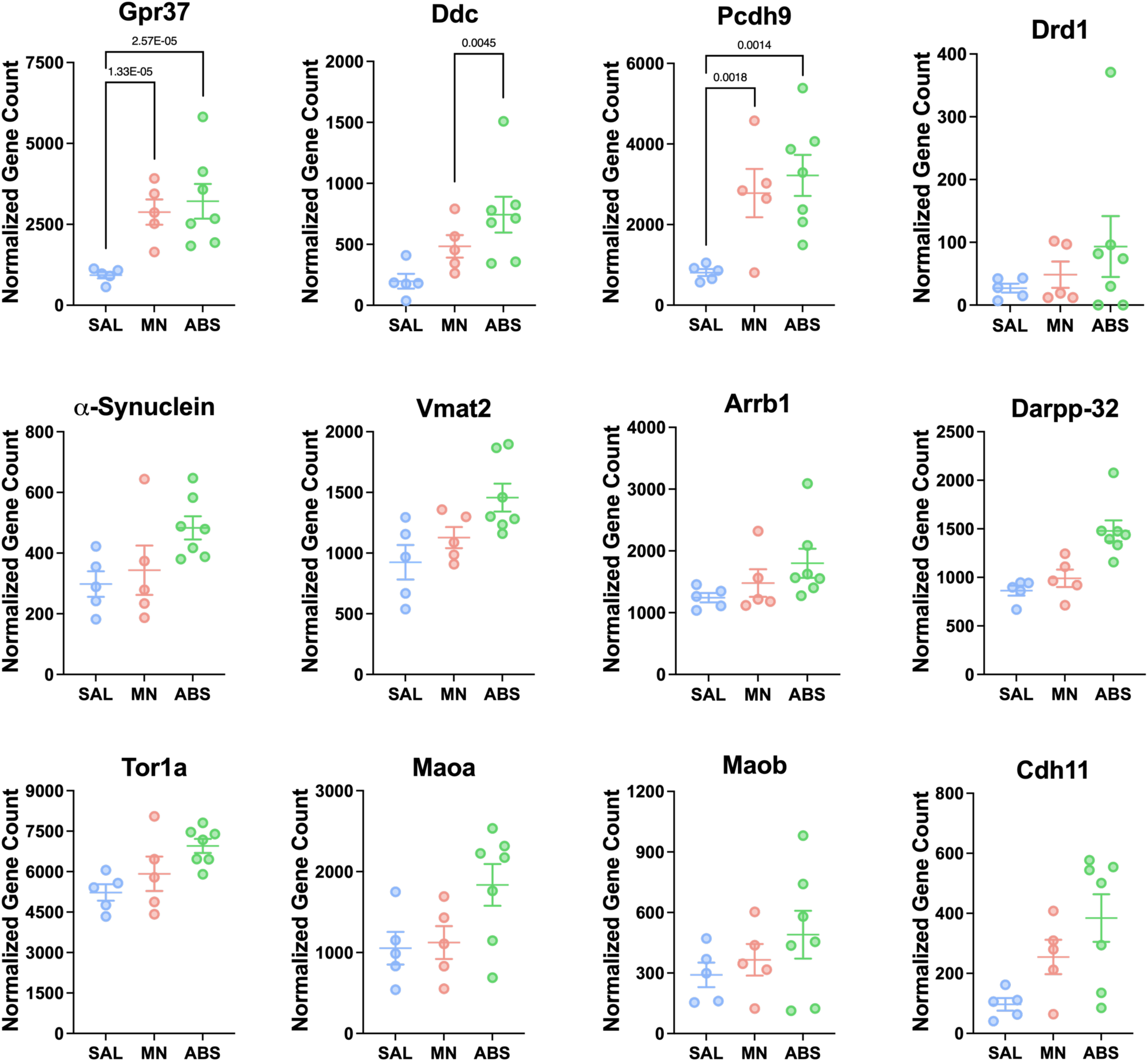
Dopamine signaling-related genes. Normalized counts of DE genes with adjusted *p-value* for each comparison. Significance shown reflects pairwise comparison results from DESeq2.

**Supplementary Figure 9.**
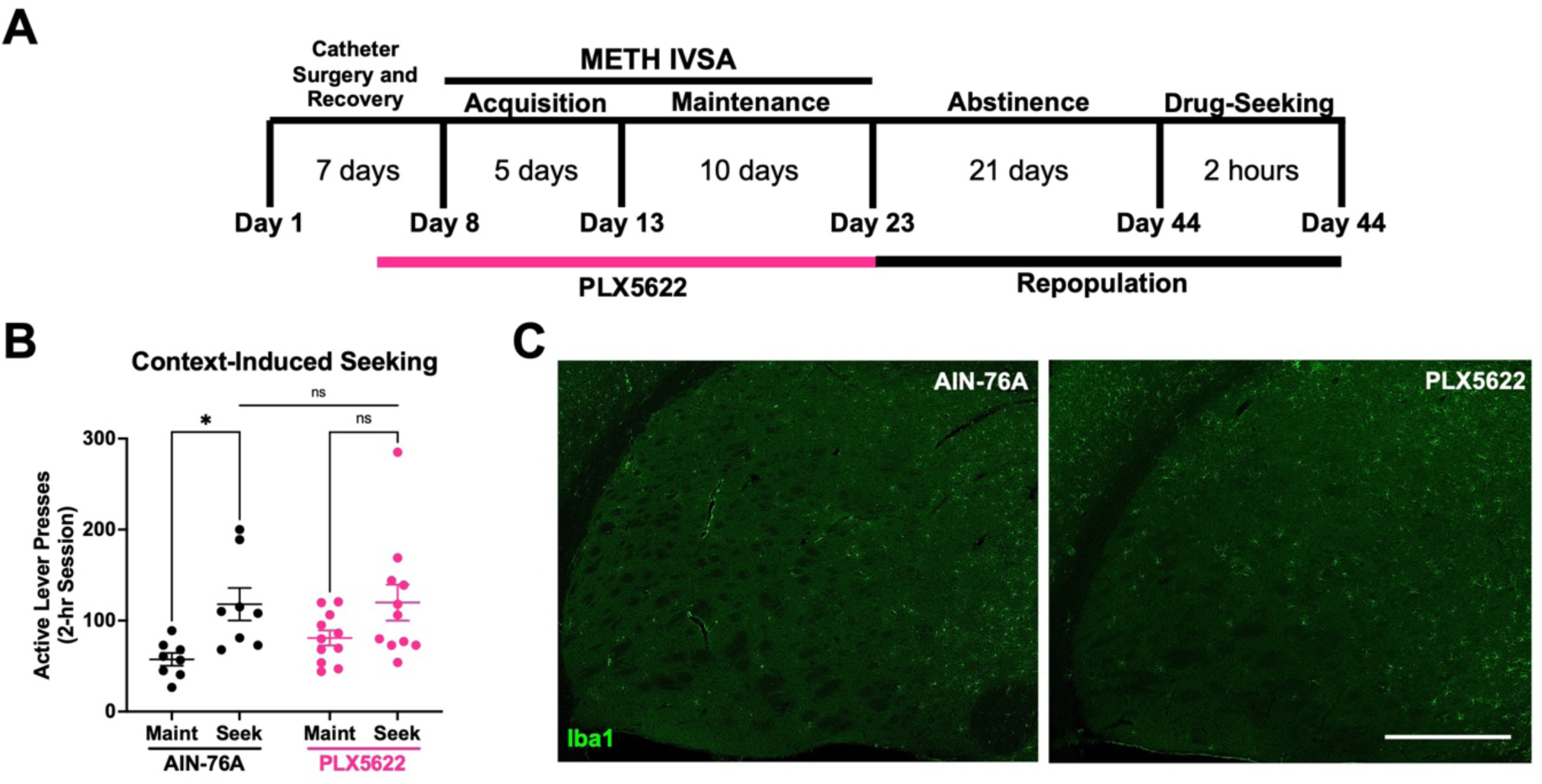
Repopulation of microglia prevents context-induced drug-seeking. Mice were treated with PLX5622 for the duration of METH IVSA before being returned to control chow (AIN-76A) for the duration of abstinence. **A)** Experimental timeline. **B)** Active lever presses for Maintenance (**Maint**: average final 3 days) and Drug-Seeking (**Seek**) of AIN-76A and PLX5622. Two-way RM ANOVA with Bonferroni post-hoc test (AIN-76A Maint vs Seek, * *p* < .05; PLX5622 Maint vs Seek, *p* = .144; AIN-76A vs PLX5622 Seek, *p* = .392). **C)** Representative fluorescent images of Iba1^+^ microglia (green) in the dorsal striatum from AIN-76A and PLX5622 treated mice. AIN-76A (n = 8), PLX5622 (n = 11). Data are represented as mean ± SEM. Scale bar = 470 μm.

